# *Mycobacterium tuberculosis* SufR Responds to Nitric oxide via its 4Fe-4S cluster and Regulates Fe-S cluster Biogenesis for Persistence in Mice

**DOI:** 10.1101/2020.08.10.245365

**Authors:** Kushi Anand, Ashutosh Tripathi, Kaustubh Shukla, Nitish Malhotra, Anil Kumar Jamithireddy, Rajiv Kumar Jha, Susmit Narayan Chaudhury, Raju S Rajmani, Arati Ramesh, Valakunja Nagaraja, Balasubramanian Gopal, Ganesh Nagaraju, Aswin Sai Narain Seshayee, Amit Singh

**Author notes:** **Correspondence** Amit Singh, Ph.D Associate Professor Wellcome Trust-India Alliance Senior Fellow Department of Microbiology and Cell Biology (MCBL) Centre for Infectious Disease Research (CIDR) Indian Institute of Science (IISc) Bangalore-12.

## Abstract

The persistence of *Mycobacterium tuberculosis* (*Mtb*) is a major problem in managing tuberculosis (TB). Host-generated nitric oxide (NO) is perceived as one of the signals by *Mtb* to reprogram metabolism and respiration for persistence. However, the mechanisms involved in NO sensing and reorganizing *Mtb*’s physiology are not fully understood. Since NO damages Fe-S clusters of essential enzymes, the mechanism(s) involved in regulating iron-sulfur (Fe-S) cluster biogenesis could help *Mtb* persist in host tissues. Here, we show that a transcription factor SufR (Rv1460) senses NO via its 4Fe-4S cluster and promotes persistence of *Mtb* by mobilizing the Fe-S cluster biogenesis system; *suf* operon (*Rv1460-Rv1466*). Analysis of anaerobically purified SufR by UV-visible spectroscopy, circular dichroism, and iron-sulfide estimation confirms the presence of a 4Fe-4S cluster. Atmospheric O_2_ and H_2_O_2_ gradually degrade the 4Fe-4S cluster of SufR. Furthermore, electron paramagnetic resonance (EPR) analysis demonstrates that NO directly targets SufR 4Fe-4S cluster by forming a protein-bound dinitrosyl-iron-dithiol complex. DNase I footprinting, gel-shift, and *in vitro* transcription assays confirm that SufR directly regulates the expression of the *suf* operon in response to NO. Consistent with this, RNA-sequencing of *Mtb*Δ*sufR* demonstrates deregulation of the *suf* operon under NO stress. Strikingly, NO inflicted irreversible damage upon Fe-S clusters to exhaust respiratory and redox buffering capacity of *Mtb* Δ*sufR*. Lastly, *Mtb*Δ *sufR* failed to recover from a NO-induced non-growing state and displayed persistence defect inside immune-activated macrophages and murine lungs in a NO-dependent manner. Data suggest that SufR is a sensor of NO that supports persistence by reprogramming Fe-S cluster metabolism and bioenergetics.

**Highlights:**

i. *Mycobacterium tuberculosis* (*Mtb*) induces the expression of *suf* operon for Fe-S cluster biogenesis in response to nitric oxide (NO).
ii. We found that a transcription factor SufR senses NO via its 4Fe-4S cluster and regulates the expression of the *suf* operon for Fe-S cluster biogenesis.
iii. SufR-regulated Fe-S cluster biogenesis confers respiratory and redox features that promote recovery of *Mtb* from NO stress.
iv. SufR activity is required to support the NO-dependent persistence of *Mtb* in macrophages and mice.

## Introduction

About 90% people infected with *Mycobacterium tuberculosis* (*Mtb*) remain asymptomatic for tuberculosis (TB). This indicates that the host immunity effectively suppresses the bacterial replication without eradicating the pathogen. The ability of the host to produce nitric oxide (NO) by an inducible nitric oxide synthase (iNOS) is known to modulate immunity [1] and microbial physiology [2], thereby controlling diverse infections [3, 4] including TB [5]. Upon infection with *Mtb*, lesional macrophages in humans and macaques express functional iNOS [6, 7], and exhaled breath of TB patient contains NO [8]. Importantly, iNOS activity seems to control TB in humans [9, 10]. Mechanistically, NO inhibits respiration and arrests growth of *Mtb* [2]. The bacterial mechanisms responsible for sensing NO and mobilizing adaptation programs are poorly understood.

Previous models suggested that *Mtb* exploits a three-component system, DosR/S/T, to induce transcriptional changes, growth arrest, and a switch from aerobic to anaerobic respiration under NO stress [11, 12]. However, the NO-mediated activation of DosR regulon is transient [13] and other gases, such as carbon monoxide (CO) and oxygen (O_2_)-limitation, similarly activate the DosR pathway [14, 15]. Therefore, the mechanism that specifically coordinates changes in metabolism, growth, and respiration of *Mtb* in response to NO is not fully understood. In this context, a recent study demonstrated active degradation of several iron-sulfur (Fe-S) cluster proteins coordinating respiration, central metabolism, and amino acid biosynthesis in NO-treated *Mtb* [13]. Importantly, a seven-gene operon, the Suf system (Rv1460-Rv1466) that is likely involved in Fe-S cluster biogenesis/repair [16], showed prolonged, elevated expression in response to NO [13]. Interestingly, excluding Rv1460 (*sufR*), other genes of the *suf* operon are essential [16, 17]. Since the reaction of NO with Fe-S clusters generates a lethal dinitrosyl-iron dithiol complex (DNIC) [18], active degradation and calibrated regeneration of Fe-S clusters via the *suf* operon represents a potential adaptive defense against NO. Despite these reports, how *Mtb* senses NO and regulates Fe-S cluster biogenesis remains uncharacterized. Filling this knowledge gap is crucial for understanding the molecular underpinning of *Mtb* persistence.

Recently, the first protein of the Suf system (SufR; *Rv1460*) has been shown to coordinate a 2Fe-2S and function as a putative regulator of the *suf* operon [19]. However, the C-terminus of SufR contains five cysteine residues (C_175_-C_179_-C_192_-C_218_-C_220_), which suggests serving as a ligand for a 4Fe-4S cluster [20, 21]. The presence of a 2Fe-2S cluster was demonstrated *in vitro* that invariably results in poor Fe-S cluster incorporation [19, 22]. Also, without a functional assay (*e.g.,* DNA binding), the authenticity of the 2Fe-2S form of SufR cannot be validated. Lastly, phenotypic characterization of the *sufR* mutant remained ambiguous as one study showed the requirement of SufR for growth under standard culture conditions but not under stress (e.g., iron-limitation) [19]. In contrast, other studies reported the opposite findings [22, 23]. Several discrepancies thus exist in the previously reported characterization of SufR, and consequently, in our understanding of its physiological function. In this study, we performed biochemical, biophysical, and genetic characterization of SufR. We show that SufR contains a NO-sensitive 4Fe-4S and regulates Fe-S biogenesis, redox balance, and bioenergetics of *Mtb* to promote persistence in response to NO *in vivo*.

## Material and Methods

### Culture Conditions

The *Mtb* H37Rv, *Mtb*Δ *sufR,* and *sufR-*comp were grown in Middlebrook 7H9 broth (Becton, Dickinson and Company (BD), USA) medium supplemented with 0.2% glycerol, 0.5% BSA, 0.2% dextrose, and 0.085% NaCl (ADS) with 0.05% Tween 80 as described previously [23]. For culturing on solid medium, *Mtb* strains were cultured on 7H10/7H11 agar medium (Becton, Dickinson and Company (BD), USA) supplemented with 1x OADC (Becton, Dickinson and Company (BD), USA) and 0.2% glycerol. *E. coli* cultures were grown in LB medium (HIMEDIA, India). Whenever required, antibiotics were added to the culture medium (for *E. coli*, 100 μg/ml kanamycin (Amresco, USA) and 150 μg/ml hygromycin (Sigma-Aldrich, India); for *Mtb* strains, 50 μg/ml hygromycin).

### Generation of the *sufR-*complemented strain

Primer pair: 5’-ATCGAAGCTTGTCCGTCCCTGCCGATCTCAC-3’ and 5’- ATGCGGTACCAACGCTCTTGCTGGCCTCTG-3’ having restriction site HindIII and KpnI were used to amplify *sufR* from wild type *Mtb*. PCR was performed and the product contained the wild-type *sufR* gene, encoded by Rv1460 with the promoter region (567 bp upstream), was cloned into pCV125 (integrated plasmid). Sequence was verified and the constructed plasmids were transformed into the *Mtb*Δ *sufR* strain. Expression of complemented strain was confirmed by qRT-PCR.

### RNA Sequencing experiments

The *Mtb* strains were grown to an O.D._600_ of 0.4 and exposed to 0.5 mM diethylenetriamine-nitric oxide (DETA/NO [Sigma-Aldrich, India]) for 4 h at 37°C. The experiment was carried out with three independent biological replicates. Total RNA extraction was conducted using the FastRNA® Pro Blue Kit (MP Biomedicals, USA) in accordance with the manufacturer’s instruction and further purified using RNeasy spin columns (Qiagen, USA) as described [23]. Following purification, the RNA was quantified and assessed for purity by a 2100 Bioanalyzer (Agilent Technologies, Waldbronn, Germany). RNA samples with an RIN (RNA Integrity Number) value >8 were processed further for sequencing. Ribosomal RNA (16s and 23s rRNA) was removed by hybridization with magnetic beads-coupled oligonucleotide (MICROBExpress Kit, Life Technologies, USA) and concentration of enriched mRNA was quantified by Qubit RNA HS Assay Kit (Life Technologies, USA). RNA-seq was performed as described [16]. In brief, libraries were prepared using NEB Next Ultra Directional RNA Library Prep Kit for Illumina (New England Biolabs, USA), according to manufacturer’s instructions. The library size distribution and quality were assessed using a high sensitivity DNA Chip (Agilent Technologies, USA) and sequenced in HiSeq 2500 platform (Illumina, USA) sequencer using 1X50 bp single-end reads with 1% PhiX spike-in control.

### Differential gene expression and statistical analysis for RNA-Seq

Raw reads were obtained for *Mtb* H37Rv strain as fastq files. The reference genome sequence (.fna) and annotation (.gff) files for the same strain (accession number: NC_000962.3) were downloaded from the ncbi ftp website (“ftp.ncbi.nlm.nih.gov”). The annotation file was customized with the addition of annotations for non-coding RNAs [62]. The format of the annotation file (.gff) was changed to .bed format using an in-house python script. The raw read quality was checked using the Fast QC software (version v0.11.5; http://www.bioinformatics.babraham.ac.uk/projects/fastqc). BWA (version 0.7.12-r1039)[63] was used to index the reference genome. Reads with raw read quality >= 20 were aligned using BWA aln -q option. SAMTOOLS (version 0.1.19-96b5f2294a) [64] was used to filter out the multiply mapped reads. BEDTOOLS (version 2.25.0)[65] was used to calculate the reads count per gene using the annotation file (.bed). The normalization and differential gene expression analysis for the conditions were carried out using edgeR as mentioned previously [66]. Genes with at least 10 reads were selected for each comparative analysis. DGE analysis was done in RStudio (1.1.447) with R version 3.4.4. (http://www.rstudio.com/)

### Aconitase assay

The activity of aconitase (Acn) was measured by monitoring the disappearance of cis-aconitate at 240nm in a UV spectrophotometer (Thermo Scientific Biomat 3S, USA) as described [67]. One unit (U) of aconitase activity is defined as 1µmol cis-aconitate formed or converted per minute. Reaction mixtures (1ml) for Acn contained 25mM Tris-HCl (pH 8.0), 100 mM NaCl, and 50 µg *Mtb* cell lysates. Reactions were initiated by adding 0.15 mM cis-aconitate and monitored by following the disappearance of cis-aconitate at 240 nm after every 15 sec for total 30 min. Absorbance at 240 nm was plotted against time. Acn activity was calculated from linear portion of the curve in initial 5 min when reaction follows 0^th^ order of reaction. An extinction coefficient of 3,500 M^-1^cm^-1^ was used to calculate the rates.

### Western blot

Whole cells lysate (50 µg) was separated on 12% SDS-PAGE and then transferred onto a PVDF membrane (GE Healthcare, Piscataway, NJ, USA). Membrane were blocked in 5% (w/v) nonfat dry milk and incubated for 3 h at room temperature with primary antibody (Acn and Cbs 1:10000 dilution). After washing with 1XTBST, membranes were incubated in goat anti-rabbit IgG HRP-conjugated secondary antibody (1:10000 dilution) for 1 h. The autoradiography signals were visualized using ECL advance Western blotting detection kit (BioRad, USA).

### OCR and ECAR measurements

The *Mtb* strains adhered to the bottom of a XF cell culture microplate (Agilent technologies, USA), at 2X10^6^ bacilli per well by using Cell-Tak (a cell adhesive). OCR and ECAR were measured using Agilent XF Extracellular Flux Analyser. Assays were carried out in unbuffered 7H9 media (pH 7.35) with glucose 2 mg/ml as carbon source. Basal OCR and ECAR were measured for initial 21 min before the automatic addition of freshly prepared DETA-NO (0 mM, 0.25 mM, 0.5 mM and 1 mM) in 7H9 unbuffered media, through port A of cartridge plate. Three measurements were taken after 1 h of incubation. CCCP (Sigma-Aldrich, India) was added at 10 µM concentration to achieve maximum rate of respiration. Raw data of OCR and ECAR was CFU normalised for 2X10^6^ CFU/ well. Spare respiratory capacity, was calculated from % OCR value, by subtracting third basal reading (normalized as 100%) from first point after CCCP addition.

### CellRox Deep Red Staining and Flow Cytometry

Logarithmically growing *Mtb* strains were treated with DETA-NO (0 mM, 0.25 mM, 0.5 mM and 1 mM) and then incubated at 37 °C with shaking for 2 h. 200 µl cells were treated with CellROX® Deep Red reagent (Thermo Fisher, USA) at a final concentration of 5 μ minutes at 37°C and analysed on BD FACSVerse flow cytometer with 640/665 nm excitation and emission respectively. We collected 5000-10,000 events for each sample wherever possible.

### Animal experiments

For the chronic model of infection, 5- to 6-week-old female BALB/c, C57BL/6 and iNOS^-/-^ mice (n = 6 per group) were infected by aerosol with approximately 100 bacilli per mouse with the *Mtb* strains using a Madison chamber aerosol generation. At indicated times post infection, mice were euthanized, and the lungs were harvested for bacillary load, tissue histopathology analysis, and pathological scoring as described [23]. The remaining tissue samples from each mouse were homogenized and bacillary load was quantified by plating serial dilutions of tissue homogenates onto Middlebrook 7H11-OADC agar plates supplemented with lyophilized BBL MGIT PANTA antibiotic mixture ((polymyxin B, amphotericin B, nalidixic acid, trimethoprim, and azlocillin, as supplied by BD; USA). Colonies were observed and counted after 4 weeks of incubation at 37°C.

### Construction of *Mtb* SufT knockdown strain

For construction of *Mtb* SufT knock down strain (SufT-KD), we have followed CRISPR interference (CRISPRi) technology was utilized as described previously [68]. Anhydrotetracycline (ATc, Cayman, USA) 200 ng/ml was added for the induction of *sufT* specific–guide RNA (sgRNA) and dCas9 every 48 h. SufT-KD culture was divided equally when A_600_ reached 0.1–0.2 and cultured in the presence or absence of ATc. The dCas9 was expressed by pRH2502 from a TetR-regulated *uvtetO* promoter and sgRNA in pRH2521 under the control of a TetR-regulated *smyc* promoter (Pmyc1tetO). To deplete *sufT,* gene specific sgRNAs were designed for two regions between 98-127 bp and 171-191 bp of *sufT* and cloned in pRH2521. Depletion of *sufT* was verified by qRT-PCR and based on the significant repression of *sufT,* we chose sgRNA targeting *sufT* region between 98-127 bp for further study. For qRT-PCR analysis, total RNA was extracted from SufT-KD and control strains after 24 h of ATc treatment. After DNase treatment, total 600 ng of RNA was used for cDNA synthesis by using Random hexamer oligonucleotide primer (iScript Select cDNA Synthesis Kit, BioRad, USA). Gene specific primers (Table S2) and iQ SYBER Green Supermix (BioRad, USA) were used for RT-PCR (StepOne Plus, Thermo, USA). Gene expression was normalized to *Mtb* 16S rRNA expression level.

### NO exposure and recovery

Logarithmically grown *Mtb* strains (OD600 of 0.8) were diluted to an OD600 of 0.1 and exposed to six doses of 100 μM DETA-NO, once every 6 h. Recovery from exposure to DETA-NO was monitored by recording the OD600 after each addition of DETA-NO, as well as every 24 h after addition. Two biological replicates were used for each strain. For single dose experiments, different doses of DETA-NO (0.5 mM, 1.25 mM, 2.5 mM and 1mM) were added. After 4 and 24 h cells were harvested, washed with 1X PBS and plated on ADS-7H11 plates. Colonies were counted after 3–4 weeks of incubation at 37°C.

### Cell line experiments

RAW264.7 murine macrophage cell line was activated by treatment with IFNγ (100U/mL, Invitrogen, USA) for 12 h before infection and LPS (100 ng/mL, Sigma-Aldrich, India) for 2 h before infection. Activated RAW264.7 macrophages were infected with wt *Mtb*, *Mtb*Δ*sufR*, and *sufR-Comp* strains at multiplicity of infection (MOI) 2 for 4 h, followed by washing thoroughly to remove extracellular bacteria with warm DMEM medium and suspended in the same containing 10% FBS. For CFU determination, macrophages were lysed using 0.06 % SDS-7H9 medium diluted in PBS/Tween and plated on OADC-7H11 at indicated time points. Colonies were counted after 3–4 weeks of incubation at 37°C.

### Purification of SufR under anaerobic conditions

The entire ORF of *Mtb sufR* (Rv1460) was PCR-amplified using gene-specific oligonucleotides (pET28a*sufR*F and pET28a*sufR*R; Table S2), digested with *Nde*I-*Hind*III, and ligated into similarly digested His-tag-based expression vector, pET28a (TAKARA BIO, Clontech Laboratories, CA, USA) to generate pET28a:SufR. A N-terminal histidine-tagged SufR was overexpressed in *E. coli* BL21 λDE3 by 0.6 mM IPTG [Isopropyl β-d-1-thiogalactopyranoside; MP Biomedicals, USA (60 min, 30 °C)]. To facilitate Fe-S cluster formation, cultures were incubated on ice for 18 min prior to induction and were supplemented with 300 μM ferric ammonium citrate and 75 μM L-methionine (Amresco, USA) and purified as described [24]. Purification was performed under strict anaerobic conditions inside an anaerobic glove box (PLas-Labs, Lansing, MI, USA) maintaining ≈2.0 ppm O2 by volume, and buffers and solutions were appropriately deoxygenated. The *sufR* gene on pET28a-SufR was mutated using oligonucleotide-based site-directed mutagenesis approach to create individual cysteine to alanine substitutions. After the PCR, *Dpn*I was added into the reaction mixture to digest the wild-type plasmid that was used as the template. The reaction mixture containing the mutated *sufR* gene was used to transform *E. coli* BL21 λDE3. Sequences of primers used to create mutations are shown in (Table S2). Resulting clones were verified by sequencing, and the mutant Cys variants of the wt SufR were purified as described earlier. In order to generate apo-SufR, the holo-SufR was incubated with EDTA (Ethylenediaminetetraacetic acid, Sigma-Aldrich, India) and potassium ferricyanide in a molar ratio of protein: EDTA: ferricyanide in 1:50:20 at 25°C and incubated for 20-30 min till the extensive loss of color. The solution was passed through PD10 desalting column and stored at -80 °C.

### UV-Vis, CD analysis, and gel filtration of SufR

The UV-visible absorption spectroscopy was carried out in a Thermo scientific spectrophotometer (Thermo scientific, USA) at 25°C. Absorption spectra of SufR WT (native/holo) and mutants were recorded immediately as the elution fractions were collected during the purification. In order to study the effect of air oxidation, on Fe-S cluster stability, freshly purified holo-SufR was transferred to an anaerobic quartz cuvette, exposed to air by opening the cap and mixing by pipetting for 2 min. The cuvette was then sealed and monitored by UV-visible spectroscopy over time (Thermo scientific, USA). To study the effect of NO, DTH and H2O2 on [4Fe-4S] cluster of SufR, the absorption spectra of freshly purified protein were recorded at indicated concentration and different time intervals in anaerobic quartz cuvette. CD measurements were conducted in a Jasco J-715 spectropolarimeter (Jasco, USA). Far-UV spectra were measured from 190 to 250 nm range and near-UV spectra from 300 nm to 650 nm range. Protein concentration used for the far-UV CD measurements was 10-20 μM and for the near-UV measurements was 140-150 μM. Cells of 1.0 cm path length were used for the measurements of the far- and near-UV spectra, respectively. Three repeat scans were obtained for each sample. The averaged baseline spectrum was subtracted from the averaged sample spectrum. The protein was dissolved in 5 mM phosphate buffer pH 7.4 and 150 mM NaCl. Results are expressed as molar ellipticity [θ] (deg cm2 dmol-1), calculated from the following formula [θ]λ=θ/[c]□l□10□n, where θ is the measured ellipticity in degrees at wavelength λ, c is the protein concentration in mgml-1, l is the light path length in centimeters and n is the number of amino acids. CD Pro software was used to analyze the data.

To determine the molecular mass of apo- and holo-SufR protein, analytical size-exclusion chromatography experiments were performed with Superdex 200 increase, 10/300 GL analytical column (GE Healthcare Life Sciences, USA). The column was pre-equilibrated and eluted with the running buffer (10 mM phosphate buffer, 10% glycerol and 100 mM NaCl at pH 7.4) at a constant flow rate of 0.5 ml/min. Molecular mass of the proteins was determined by using gel filtration molecular mass standard (Carbonic Anhydrase, Albumin, Alcohol Dehydrogenase, β-Amylase, Apoferritin and Thyroglobulin). Ve/Vo was plotted as a function of log10Mr of the standard protein where Ve is the elution volume of the protein, Vo is the void volume of the column and Mr is the molecular weight of the particular protein. Blue dextran was used to determine the void volume (Vo). Running buffers were purged with nitrogen gas and were degassed thoroughly to remove any dissolved atmospheric oxygen. The experiments were conducted under anaerobic conditions.

### EPR spectroscopy of SufR

For EPR spectroscopy, holo SufR was treated inside the anaerobic glove box with aliquots of freshly prepared proline NONOate (Cayman Chemicals, Ann Arbor, MI, USA). Aliquots of SufR were placed in an anaerobic cuvette and titrated by injection with aliquots of a 2.5 mM stock solution of proline NONOate (Cayman chemicals, USA). Samples were then transferred to EPR tubes and immediately frozen in liquid nitrogen. EPR characteristics of NO-treated SufR were analyzed by subjecting the samples to continuous-wave spectrometers at liquid nitrogen temperature as described previously [25]. NO-treated SufR was analyzed by continuous-wave EPR on a perpendicular mode X-band EPR spectrometer operating at 100-kHz modulation frequency and equipped with liquid nitrogen cryostat and a dual mode X-band cavity (JES200 ESR spectrometer, JEOL, USA). Field calibration was done by using a standard NMR G meter. The background signal from the buffer was subtracted from the spectra.

### Chemical analysis of iron and sulfide

The total iron content and acid-labile sulfide content of holo-SufR, was measured using a previously described procedure [69]. For iron estimation, freshly purified holo-SufR (0.1 mL) was heated at 95°C degree for 30 min after treatment with 22% HNO_3_ (0.1 mL). Samples were cooled to ambient temperature followed by addition of 0.6 mL ammonium acetate (7.5 % w/v), 0.1 mL freshly prepared ascorbic acid (12.5% w/v) and 0.1mL ferene (10 mM). The concentration of iron present in the protein was determined by measuring the absorbance of the product, iron-ferene complex at 593 nm, which was compared with a standard curve prepared from dilutions of freshly prepared Fe(III) solution in the range of 0-200 µM. To measure acid-labile sulphide content, freshly prepared Na_2_S.9H_2_O solution was used to prepared standard solution in the range of 52-260 µM of S^2-^. Protein sample/standard (200 µL) was mixed with 0.6 mL of zinc acetate (1% w/v) followed by addition of 50 µL of NaOH (12% w/v). After incubation of 15 min, 0.15 mL of N, N-dimethyl-p-phenylenediamine dihydrochloride (0.1% w/v dissolved in 5 M HCl) and 0.15 mL of freshly prepared 10 mM FeCl_3_ (dissolved in 1 M HCl) was added. The reaction mixture was further incubated for 30 min at room temperature and the absorbance of the product, methylene blue, was measured at 670 nm. For iron and sulfide estimation three independent preparations of holo-SufR were analyzed to ensure the reproducibility and three dilution of each sample were considered.

### Electrophoretic mobility shift assay (EMSA)

For EMSA, promoter fragment of *sufR* (170 bp upstream of ATG) and *blaC* (200 bp upstream of ATG) were PCR amplified from the *Mtb* genome. The 5’ end was labeled by [Y-32_P_]-ATP using T4 polynucleotide kinase (MBI Fermentas, USA) as per the manufacturer’s instructions. The labeled oligonucleotides were passed through a 1 mL Tris-EDTA, pH 7.5 equilibrated Sephadex G-50 column and elute was vacuum dried. Blunt ended duplex DNA was prepared by annealing radiolabelled and complementary cold oligonucleotides in a 1:2 molar ratio in 50 μL reaction system containing TE and 1X saline-sodium citrate buffer (3 M sodium chloride and 0.3 M sodium citrate). The reaction mixture was heated to 95 °C for 5 min and then cooled in a thermal cycler. Annealed mixtures were resolved by 10% native PAGE in 1X Tris-borate EDTA (TBE). Assembled DNA substrates were visualized on an X-ray film and substrates were subsequently purified by excising the respective desired bands. DNA substrates were eluted by incubating corresponding gel pieces in TE at 4°C for 6 h. Binding reactions were performed in binding buffer (25mM Tris-HCl, 1mM DTT, 0.1 mg/ml BSA, 5 mM MgCl_2_; pH 7.4) for 30 min at 4°C and 6% polyacrylamide gel was used to resolve protein-DNA complexes in an anaerobic chamber by electrophoresis at constant voltage (50 V). 0.5 nM ^32^P-labeled DNA substrates were incubated with indicated concentrations of various forms of SufR (apo, holo, H_2_O_2_-, and NO-treated). For competition with unlabeled DNA, fragments of *sufR* and *blaC* (∼200–250 bp upstream of translational start codon) were PCR amplified from the *Mtb* genome and used in various amounts to outcompete binding of holo-SufR to ^32^P-labelled DNA fragments. Gels were exposed to auto radiographic film and visualized by a phosphor imager (Typhoon FLA-9000, GE Healthcare Life Sciences, USA). All the above reactions and separations were performed under anaerobic condition inside glove box (Plas-Labs, Lansing, MI, USA).

### *In vitro* transcription assays

The DNA templates (180 bp) including the *sufR* promoter regions were PCR amplified using primers P*sufR* F1/P*sufR* R1 (Table S2). The amplicons (50 nM) were pre-incubated with different concentration of holo- and apo-SufR in the transcription buffer (50 mM Tris HCl, (pH 8.0), 10 mM magnesium acetate, 100 mM EDTA, 100 mM DTT, 50 mM KCl, 50 mg/ml BSA, and 5% glycerol) for 30 min at room temperature. Single-round transcription reactions were initiated with the addition of 100 nM *Mtb* RNAP-σA holo enzyme, 100 μM NTPs, 1 μCi [α-32P]-UTP, 50 μg ml−1 heparin and incubated at 37°C for 20 min. The reactions were terminated by addition of 2X formamide dye (95% formamide, 0.025% (w/v) bromophenol blue, 0.025% (w/v) xylene cyanol, 5 mM EDTA and 0.025% SDS and 8 M urea) and heated at 95 _C for 5 min followed by snap chilling in ice for 2 min. The transcripts were resolved on an 8% TBE400 urea-PAGE gel. To study the effect of NO, transcription assays were performed as mention above after treating the holo-SufR with proline NONOate for 5 min followed by purification. All the treatments and the reactions were performed inside anaerobic glove box under anoxic condition.

### DNase I Footprinting

5 nM of ^32^P-labeled DNA substrate was incubated with increasing concentration of purified holo-SufR in binding buffer (25 mM Tris-HCl (pH 7.5), 1 mM DTT, 100 μg/ml BSA, 5 mM MgCl2, 5 mM CaCl2), and samples were incubated for 30 min at 4°C inside glove box under anaerobic condition. Reactions were initiated by the addition of DNase I to a final concentration of 0.05 units and incubated for 2 min at room temperature. The reactions were terminated by the addition of 150 µL of TE (pH 7.5) followed by incubation at 75°C for 15 mins to deactivate DNase I enzyme. The sample was further subjected to vacuum evaporation and the pellet thus obtained was re-suspended in loading dye (80% (v/v) formamide, 0.1% (v/v) BPB, and 0.1% (v/v) xylene cyanol) and analyzed on a denaturing 15% polyacrylamide gel containing 7M urea. The gel was dried, and the bands were visualized with a Typhoon FLA-9000 phosphor imager (GE Healthcare Life Sciences, USA). Maxam and Gilbert A+G ladder was prepared as described previously [70]. A custom python script was developed to search for the SufR binding site and find its relative positions with respect to genes across the genome. A sequence of 15 nt length was searched in the region 500 bp upstream of the start codon and considered as a potential SufR binding site if it shares 80% identity with ’ACACT’ or ’TGTGA’ at the ends.

### Ethics

Animal experimentation: This study was carried out in strict accordance with the guidelines provided by the Committee for the Purpose of Control and Supervision on Experiments on Animals (CPCSEA), Government of India. The protocol of animal experiment was approved by animal ethical Committee on the Ethics of Animal Experiments, Indian Institute of science (IISc), Bangalore, India (Approval number: CAF/Ethics/544/2017). All efforts were made to minimize the suffering.

### Statistical Analysis

All data were graphed and analyzed with Prism v8.0 (GraphPad) unless otherwise stated. Statistical analyses were performed using Student’s t-test (two-tailed). Where comparison of multiple groups was made either one-way or two-way ANOVA with Bonferroni multiple comparisons was performed. Differences with a p value of <_0.05 were considered significant. Statistical significance for RNA-seq was calculated using the QL F-test followed by Benjamini436 hochberg method of multiple testing correction.

### Miscellaneous

The molar extinction coefficient per [4Fe-4S]^2+^ cluster (413=16200 M^-1^ cm^-1^) was determined The molar_extinction coefficient per [4Fe-4S]2+ cluster (ε413=16200 M-1 cm-1) was determined from *in vitro* reconstituted protein. Protein concentration of SufR throughout the study was calculated using A280 reading. Anti-SufR polyclonal antibody was used to monitor the expression and purification of wt SufR and mutant proteins by Western blot. Samples (5 mg of protein per slot) were resolved by 12% reducing SDS-PAGE and transferred to nitrocellulose membrane. Blots were probed with primary and secondary (horseradish peroxidase-conjugated anti-rabbit IgG) antibodies and processed.

### Data availability

The RNA-sequencing data presented in the manuscript are deposited into the NCBI Gene Expression Omnibus (GEO) (https://www.ncbi.nlm.nih.gov/geo/query/acc.cgi?acc=GSE154169 with the accession number GSE154169 and reviewer token kbcdmqecrlotzgp.

## Results and Discussion

### *Mtb* SufR contains a 4Fe-4S cluster

To understand the role of *Mtb* SufR in sensing NO and regulating Fe-S cluster homeostasis, we first examined the biochemical and biophysical characteristics of the SufR Fe-S cluster. We anaerobically purified histidine-tagged SufR from *E. coli* cultured in growth conditions optimized for the maximum incorporation of Fe-S clusters *in vivo* [24]. As isolated, native SufR (holo-SufR) displayed a characteristic straw brown color, with an absorption maximum indicative of a 4Fe-4S cluster at 413 nm (molar absorption coefficient [ε] at 413 = 16200 M^-1^ cm^-1^) (Fig 1A and Fig. S1 A-C) [25]. Treatment of holo-SufR with a one-electron donor, sodium dithionite (DTH), caused partial bleaching of brown color and loss of absorbance at 413 nm, consistent with the presence of a redox-responsive 4Fe-4S cluster [25] (Fig. 1A). Gel filtration performed under anaerobic conditions suggested a molecular mass of ∼ 54 kDa for native SufR, consistent with a dimer. Clusterless SufR (apo-SufR) also eluted as a dimer, indicating that the Fe-S cluster does not bridge SufR monomers (Fig. 1B). Moreover, the identity of cysteine residues that coordinates the 4Fe-4S cluster was experimentally validated. The alanine mutant of two putative cysteine ligands (C_175_A_C_179_A) was sufficient to disrupt the 4Fe-4S cluster as evident from the loss of absorbance at 413 nm (Fig. 1C).

**Figure 1:**
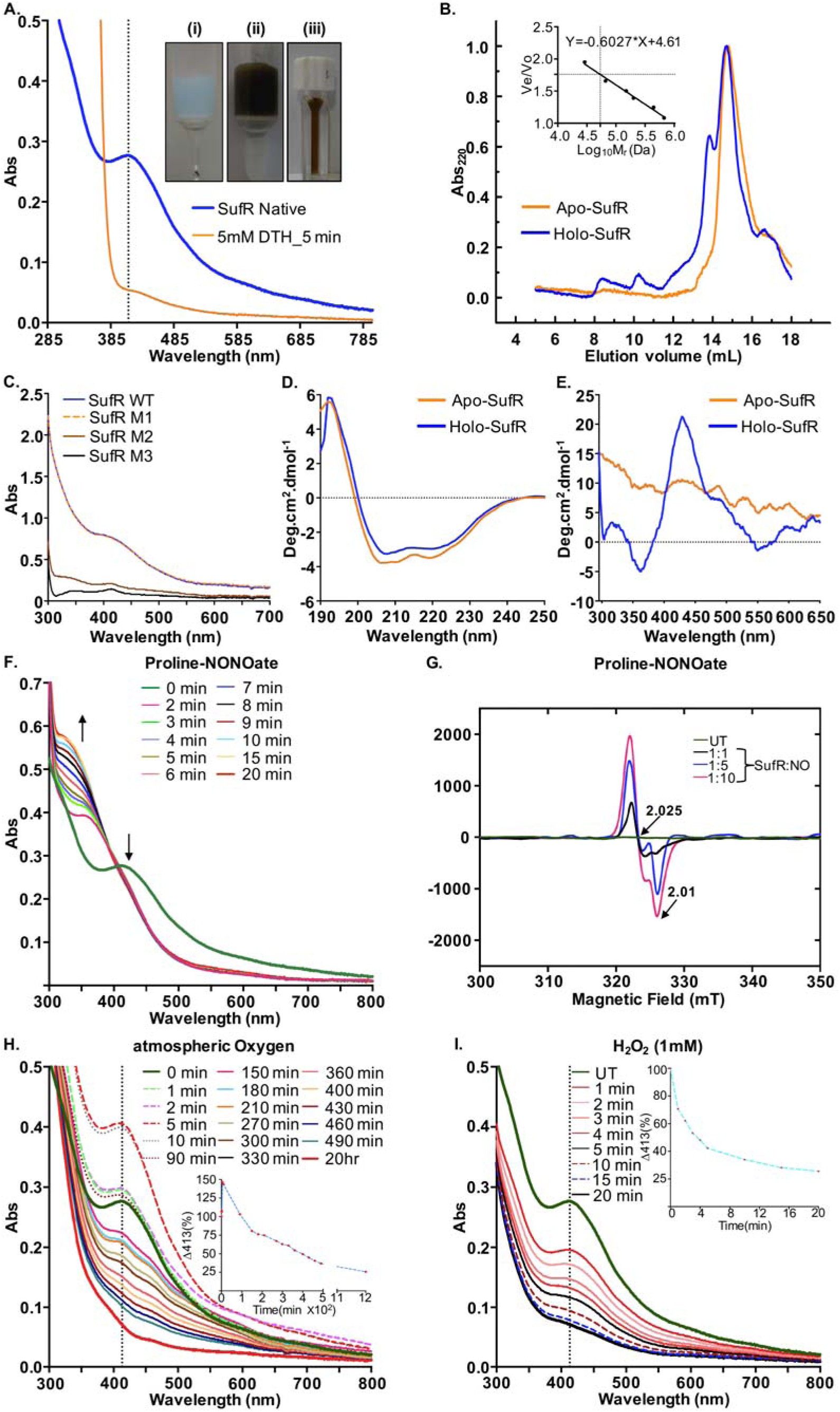
*Mtb* SufR coordinates 4Fe-4S cluster that responds to NO and H_2_O_2_. (**A**) Anaerobically-purified SufR was subjected to UV-visible spectroscopy. A clear peak at 413 nm indicates presence of a 4Fe-4S cluster. The Fe-S cluster was reduced by addition of 5 mM DTH. (*Inset)* (**i**) unbound Ni^2+^-NTA, (**ii**) brown colored SufR bound to Ni^2+^-NTA beads, and (**iii**) anaerobically purified holo-SufR. (**B**) Superdex 200 HR gel filtration chromatography of SufR. (**Inset**: standard curve of the protein standards versus their elution parameter Ve/Vo where, Ve is the elution volume and Vo is void volume). (**C**) UV-visible spectroscopy of cysteine mutants of SufR (M1-C_175_A, M2-C_175_A_C_179_A, and M3-C_175_A_C_179_A_C_192_A). (**D**) A far-UV and **(E)** Near- UV CD-spectrum of holo- and apo-SufR. (**F**) UV-visible spectra were acquired before and after addition of proline NONOate (2.5 mM). A time dependent increase in absorbance at 350 nm indicates monomeric DNIC formation. (**G**) Holo-SufR exposed to various concentration of proline NONOate (SufR:NO). EPR data were acquired using a continuous-wave EPR spectrometer at a microwave frequency of 9.667 GHz and microwave power of 2 mW at liquid nitrogen temperature. The peak at 2.025 g is consistent with the formation of monomeric DNIC. (**H**) UV-visible spectra of holo-SufR exposed to either atmospheric O_2_ for 2 min or (I) H_2_O_2_ (1 mM). (*Inset*) Rate of the 4Fe-4S cluster loss was determined by calculating the percent loss of absorbance at 413 nm upon exposure to air and H_2_O_2_.

A recent study using aerobically purified SufR, demonstrated that SufR is a monomer and coordinates a 2Fe-2S cluster upon Fe-S cluster reconstitution *in vitro* [19]. In general, Fe-S clusters are inherently unstable under aerobic conditions [26] and stabilize only under anaerobic conditions [24]. Fe-S clusters can be assembled *in vitro*, but this process has poor yield [24]. Therefore, the technique of assembling the Fe-S cluster *in vivo* followed by anaerobic purification is more likely to provide native configuration of SufR Fe-S cluster. We clarified this by analyzing native SufR using circular dichroism (CD). The far-UV CD-spectrum showed two minima at 208 and 224 nm and indicated 70% α-helical content in the secondary structure (Fig. 1D). The near-UV CD-spectrum displayed two characteristic maxima at 330 and 420 nm and two minima near 350 nm and 550 nm, as described earlier for [4Fe-4S] cluster (Fig. 1E) [24, 27] As expected, the near-UV CD-spectrum of apo-SufR did not show a [4Fe-4S] cluster (Fig. 1E). Lastly, the estimation of iron and sulfide ions in the native SufR revealed the association of 3.23-3.75 iron atoms per SufR monomer and similar amount of sulfide ions per SufR monomer (Fig. S1D). In sum, we demonstrate that *Mtb* SufR contains a 4Fe-4S cluster.

### SufR responds to NO through its 4Fe-4S cluster

We exposed native SufR to the fast-releasing NO donor; proline NONOate (T_1/2_ ∼ 1.8 s, pH 7.4) and subjected it to UV-visible spectroscopy. Treatment with proline NONOate gradually reduced absorbance at 413 nm and formed a new chromophoric feature at ∼350 nm over time with a clear isosbestic point (Fig. 1F). These spectral features were consistent with a dinitrosyl-iron dithiol complex (DNIC), wherein the sulfide ligands of the 4Fe-4S clusters were displaced by NO to form [Fe-(NO)_2_] [28]. The DNIC formation was confirmed by subjecting NO-treated SufR to continuous-wave electron paramagnetic resonance (cw-EPR). A strong EPR signal centered at *g*=2.03 was observed in a dose-dependent manner, consistent with the formation of the monomeric DNIC complex on SufR (Fig. 1G) [25].

The *suf* operon showed induction under oxidizing stress (*e.g.,* H_2_O_2_) [29]. Therefore, we determined whether the Fe-S cluster of SufR is responsive to atmospheric O_2_ and H_2_O_2_. Exposure to atmospheric O_2_ resulted in a slow decline in absorbance at 413 nm (Fig. 1H). A plot of ΔA_413nm_ against time revealed the loss of ∼75% of the 4Fe-4S cluster only at 20 hours (h) post- O_2_ exposure. However, exposure of native SufR to 1 mM H_2_O_2_ resulted in the rapid loss of absorption at 413 nm (Fig. 1I). A plot of ΔΔ_413nm_ against time revealed the loss of ∼75% of the absorption at 413 nm (Fig. 1I). A plot of Δ _413nm_ 4Fe-4S cluster in 20 min. Higher concentrations of H_2_O_2_ (10 mM and 100 mM) resulted in the loss of the cluster within 1-2 min (Fig. S2A-C). Taken together, these data demonstrate that the 4Fe-4S cluster of SufR is sensitive to NO and H_2_O_2_.

### SufR binds upstream of *suf* operon promoter and represses expression

To investigate if SufR mediates Fe-S cluster biogenesis by regulating the *suf* operon expression, we first assessed the DNA-binding properties of SufR. We carried out electrophoretic mobility shift assays (EMSA) using a radioactively (^32^P)-labeled 170 bp DNA fragment encompassing the promoter region of *suf* operon [19]. We used four different forms of SufR; holo-SufR, apo-SufR, NO-treated SufR, and H_2_O_2_-treated SufR. Holo-SufR bound to the *suf* promoter region, whereas apo-SufR, NO-treated SufR, and H_2_O_2_-treated SufR showed no DNA binding (Fig. 2A-C). DNA binding was outcompeted by 50-100-fold excess of unlabeled *suf* promoter DNA, but not by an unrelated promoter fragment (*blaC*) (Fig. 2D). These findings suggest that *Mtb* SufR is an NO- and H_2_O_2_-responsive, sequence-specific DNA-binding protein.

**Figure 2:**
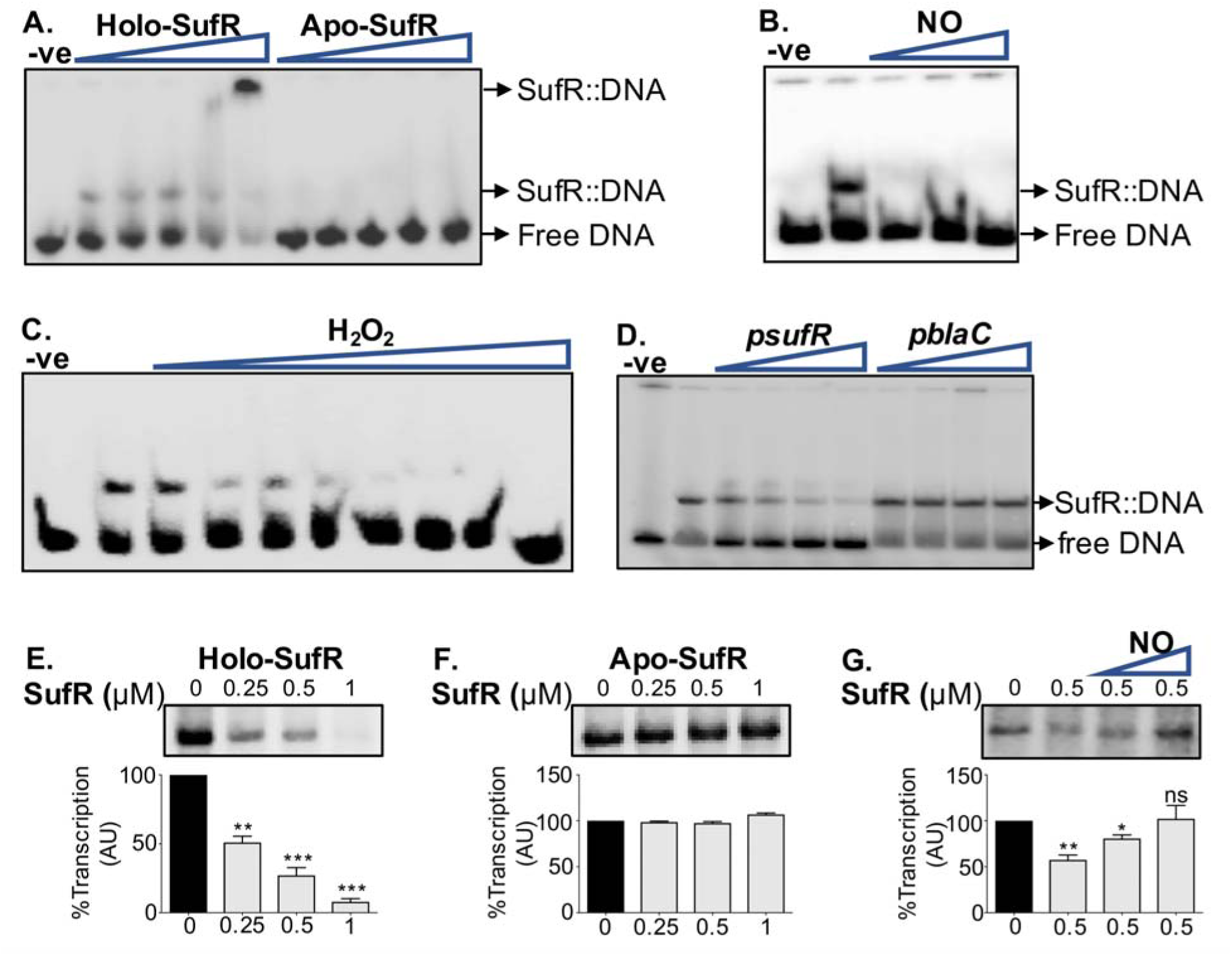
DNA binding activity of SufR. (**A**) Binding of SufR to ^32^P-labelled *suf* promoter (0.5 nM). The concentrations of SufR were 0.125 µM, 0.25 µM, 0.5 µM, 1 µM, and 2 µM. Holo-SufR (0.5 µM) was pretreated with (**B**) proline NONOate (NO; 2.5-, 5-, and 10-fold molar excess) or (**C**) H_2_O_2_ (25, 50, 75, 100, 250, 500, 1000 and 5000 nM) and were incubated with ^32^P-labelled *suf* promoter. (**D**) Holo-SufR (0.5 µM) was incubated with ^32^P-labelled *suf* promoter and DNA binding was competed out using 12.5-, 25-, 50-, and 100-fold molar excess of unlabelled competitor promoter of *suf* operon and *blaC*. (**E**) Effect of SufR on transcription from *suf* promoter *in vitro*. Increasing concentrations of holo-SufR inhibit transcription. (**F**) Apo-SufR was completely ineffective in inhibiting transcription. (**G**) Holo-SufR (0.50 µM) pre-treated with proline NONOate (NO; 2.5- and 5-fold molar excess) stimulated transcription from *suf* promoter. The graph below indicates densitometry analysis of the *suf* transcript (n=2). One-way analysis of variance (ANOVA) with Bonferroni’s post hoc was used to determine statistical significance. ‘ns’ non-significant ‘*’ p<0.05 ‘**’ p<0.01 ‘***’ p<0.001.

We next performed *in vitro* transcription using *Mtb* RNA polymerase as described [30]. A single round of transcription of the *suf* promoter fragment (180 bp [-73 to + 107 bp]) produced a single transcript of 107 nucleotides, consistent with the leaderless transcription of the *suf* operon [31]. Addition of holo-SufR repressed transcription from the *suf* promoter in a dose-dependent manner, whereas apo-SufR did not (Fig. 2E-F). Importantly, the treatment of holo-SufR with proline NONOate (NO) reversed the repression of the *suf* promoter (Fig. 2G). These results indicate that holo-SufR binds to the *suf* promoter and represses expression, whereas NO-damaged or clusterless apo-SufR lacks DNA-binding and transcription repression. The data suggest that occupancy of Fe-S cluster on SufR serves as an indicator of Fe-S cluster-sufficient or -deficient conditions. During low Fe-S cluster demand, holo-SufR represses the Suf system to limit excessive assembly of Fe-S clusters. In contrast, loss or damage of the Fe-S cluster in SufR (apo-SufR) signals heightened demand for Fe-S clusters that results in the de-repression of the *suf* operon and mobilization of Fe-S cluster assembly. In support of this, we depleted an essential gene of the *suf* operon (Rv1466; *sufT*) involved in Fe-S cluster maturation [32] using CRISPR interference (CRISPRi-*sufT*). The depletion of SufT is expected to perturb Fe-S cluster biogenesis and increase the pool of apo-SufR, which could result in the de-repression of the *suf* operon in *Mtb*. Consistent with this, expression of the *suf* operon was induced in CRISPRi-*sufT* as compared to wild-type *Mtb*, indicating that *Mtb* de-represses the *suf* operon in response to abnormal Fe-S cluster biogenesis caused by SufT depletion (Fig. S3).

Lastly, we performed DNase I footprinting to identify the binding site of holo-SufR on the promoter region of the *suf* operon (Fig. 3A). A clear region of protection from DNase I digestion was evident with increasing molar ratios of SufR:DNA (Fig. 3A-B). The protected region contains a perfect inverted repeat (ACACT-N_5_-TGTGA) separated by 5 bp (Fig. 3A-B). This inverted repeat forms a part of a larger inverted repeat (ATTTTGTCACACT-N_5_-TGTGAAAAT). Consistent with the footprinting data, mutations in the inverted repeat completely abolished binding, thus confirming that the SufR binds to the palindrome ACACT-N_5_-TGTGA (Fig. 3 C-E). Altogether, using multiple techniques, we confirmed that SufR functions as a NO-sensitive DNA-binding transcription factor in *Mtb*.

**Figure 3:**
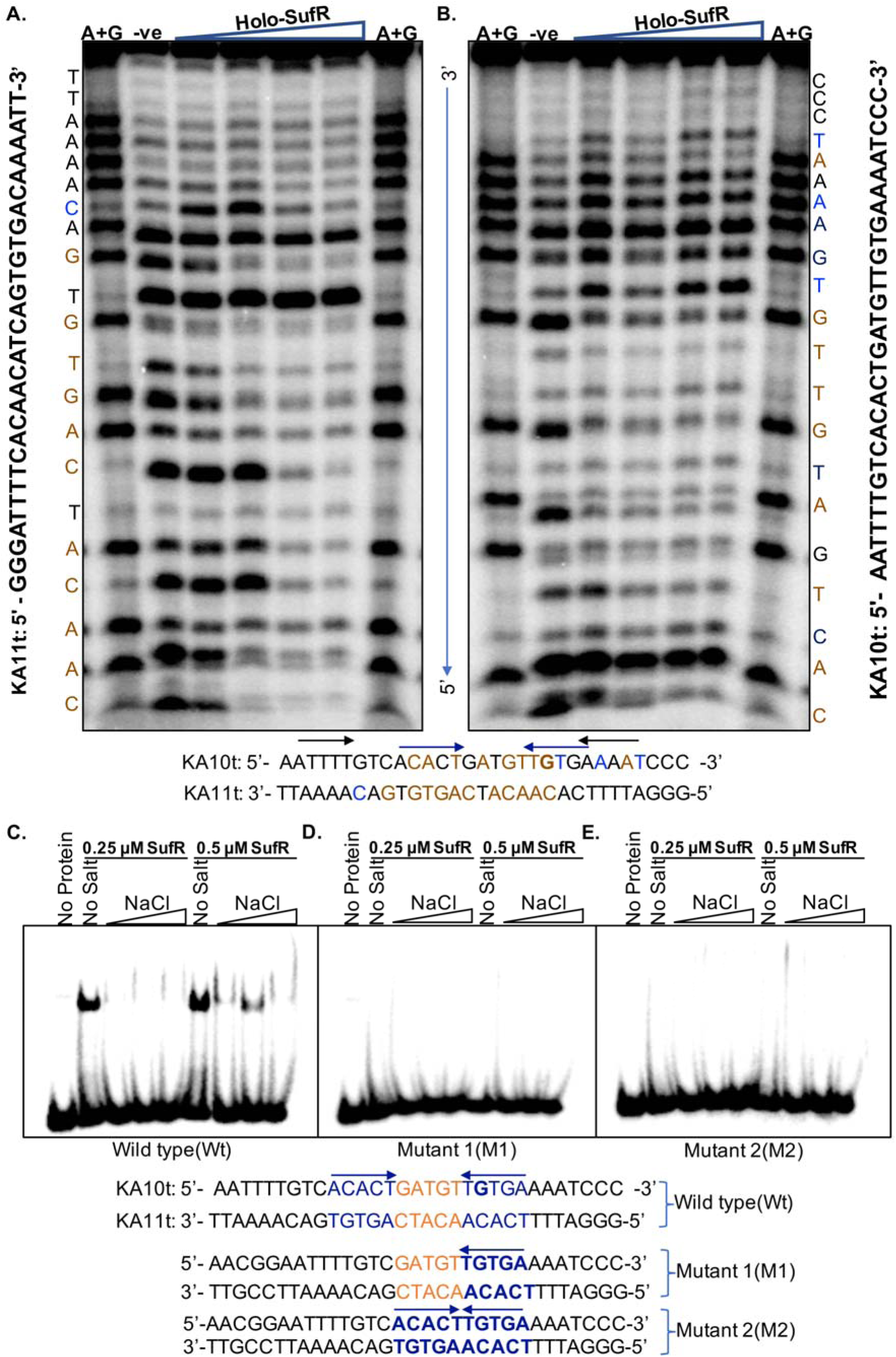
DNase I footprint of holo-SufR bound to the *suf* promoter. Sequence upstream of the *suf* operon (KA10t/KA11t) were used for DNase I footprinting. Bold letter G represents transcription start site (TSS) for the *suf* operon. (**A**) ^32^P-KA10t was annealed with cold KA11t. (**B**) ^32^P-KA11t was annealed with cold KA10t to examine footprint in the reverse strand. 5 nM^32^P-labeled dsDNA was incubated either in absence or presence of holo-SufR (125 nM, 250 nM, 500 nM, and 1 µM) and treated with 0.05 U of DNase I for 2 min. A+G is Maxam and Gilbert sequencing ladder. DNase I protected/unprotected/hypersensitive nucleotides are marked with brown/black/blue and two inverted repeats are marked by arrows (blue and black). **Mutations in the inverted repeat completely abolished binding of holo-SufR. (C-E)** EMSA was performed with wild type (KA10t and KA11t) and oligonucleotides with mutations in the inverted repeats (KA10tm1, KA11tm1, KA10tm2 and KA11tm2) at increasing concentrations of NaCl at two different concentrations of holo-SufR (0.25 µM and 0.5 µM).

### NO regulates expression of the *suf* operon and Fe-S cluster pathways in *Mtb*

Having shown that SufR responds to NO via its 4Fe-4S cluster and directly regulates the transcription of *suf* operon promoter, we next asked if SufR coordinates *Mtb*’s adaptation under NO stress. To examine this possibility, we first investigated the role of SufR in regulating transcriptome of *Mtb* in response to NO. All genes of the *suf* operon are essential except *sufR*. [16]. Therefore, we utilized *sufR*-deficient (*Mtb*Δ*sufR)* strain of *Mtb* [23]. We generated the *sufR*553 complemented strain (*sufR-Comp*) by integrating *sufR* (Rv1460) along with its native promoter (∼500 bp upstream of *sufR*) in the genome of *Mtb*Δ*sufR.* Using qRT-PCR, we confirmed the restoration of *sufR* expression to wild type (wt) *Mtb* levels in *sufR-Comp* (Fig S4.). Since exposure to 0.5 mM of NO donor diethylenetriamine-nitric oxide (DETA-NO) arrested the growth of wt *Mtb*, *Mtb*Δ*sufR*, and *sufR-Comp* without affecting viability (Fig. S5), we performed RNA-sequencing (RNA-seq) on *Mtb* strains treated with 0.5 mM of DETA-NO for 4 hours.

Our RNA-seq data recapitulated the previously published NO-responsive transcriptome of *Mtb*, with the induction of the DosR, IdeR, Zur, and CsoR regulons, and reduced expression of RNA and protein biosynthesis pathways (fold change≥2; FDR≤0.05; Fig. S6 and Table S1A) [13]. These pathways were similarly affected in *Mtb*Δ*sufR* and *sufR-Comp* in response to NO (Fig. S6 and Table S1B-C). Since Fe-S cluster proteins are the most susceptible targets of NO [33], we found that the expression of ∼50% of genes encoding Fe-S cluster proteins was altered in wt *Mtb*, *Mtb*Δ*sufR*, and *sufR-Comp* under NO stress (Fig. 4 and Table S1D). The strong reactivity of NO towards heme iron of primary cytochrome oxidase is thought to arrest respiration [2]. However, we found that NO does not affect the expression of genes encoding primary terminal oxidase (cytochrome bc1-aa3 complex; *qcrABC*) in *Mtb* (Fig. 4). Nonetheless, downregulation of Fe-S cluster-containing NADH dehydrogenase complex I (*nuo* operon) and other genes encoding Fe-S cluster proteins involved in central metabolism (*acn, prsA*, and *udgA*) are consistent with the inhibition of primary respiration by NO. Moreover, the expression of genes associated with alternate form of respiration (*e.g.,* NADH dehydrogenase type II [*ndh*], cytochrome BD oxidase [*cydABCD*], nitrate reductase [*narH*], sulfite reductase [*sirA*], hydrogenase [*hycP*]) [34], carbon catabolism (*korAB*, *frdB, icl1,* and *pckA*), and branched chain amino acid (BCAA) biosynthesis [35] were induced by NO. (Fig. 4).

**Figure 4:**
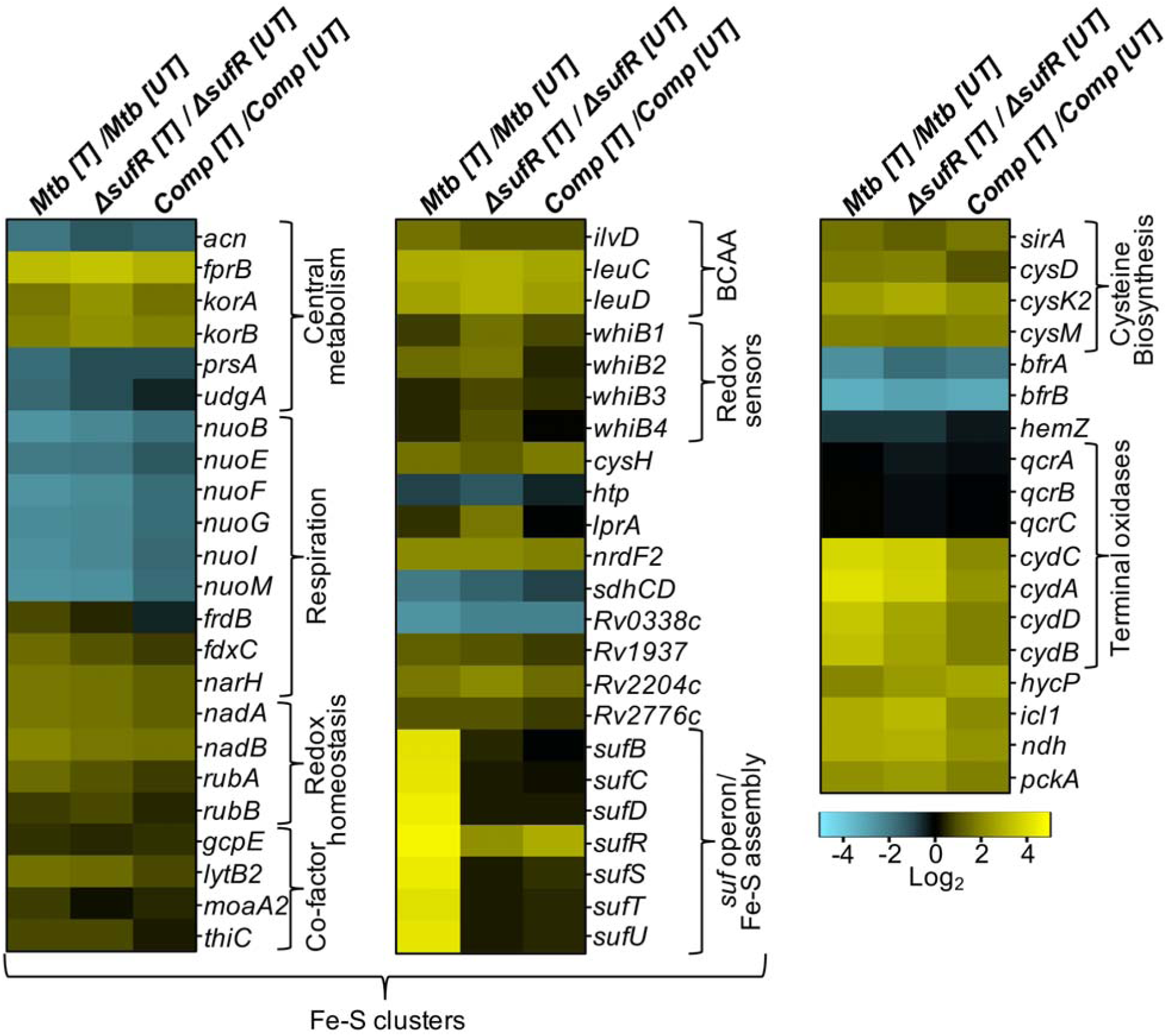
SufR mainly regulates Suf system involved in Fe-S cluster biogenesis in response to NO. Total RNA from three biological replicates of untreated (UT) and DETA-NO treated (T) wt *Mtb*, *Mtb*Δ*sufR,* and *sufR-Comp* was isolated and subjected to RNA-seq. Heat maps indicate log_2_ fold change of differentially expressed genes (DEGs) belonging to various functional categories (obtained from Mycobrowser, EPFL, Lausanne). Genes were considered differentially expressed on the basis of the false discovery rate (FDR) of ≤0.05 and absolute fold change of ≥1.5. BCCA: branched-chain amino acid.

The association between NO, respiration, and metabolism is further indicated by the upregulation of Fe-S cluster enzymes associated with the biosynthesis of respiratory cofactors such as isoprenoid (*lytB2* and *gcpE*), molybdopterin (*moaA2*), thiamin (*thiC*), quinolinate (*nadA-B*), and sulfur metabolites (*cysH*) [36–39] (Fig. 4; Table S1D). Also, NO induced the expression of transcription factors containing redox-sensitive Fe-S clusters (*e.g., whiB1, whiB2,* and *whiB3)* [25, 40, 41] (Fig. 4). We found that NO induced comparable expression changes in *Mtb* and *Mtb*Δ*sufR* (Fig. 4), indicating that SufR is not a global regulator of *Mtb*’s transcriptome under NO stress.

Upregulation of the Fe-S cluster transcriptome indicates an increased demand for Fe-S cluster biogenesis. In agreement, expression of the *suf* operon (*sufRBDCSUT*; Rv1461-Rv1466) was stimulated 15- to 25-fold by NO in *Mtb* (Fig. 4). Also, NO induces the cysteine biosynthetic machinery that supplies sulfur for Fe-S cluster biogenesis (Fig. 4). However, genes coordinating biosynthesis of Fe-binding heme (*hemZ*) and encoding Fe-storage proteins (bacterioferretin; *bfrA-B*) were downregulated (Fig. 4). These observations suggest that *Mtb* prioritizes the assembly of Fe-S clusters under NO stress. In contrast to wt *Mtb*, the *suf* operon remains basally expressed in NO-treated *Mtb*Δ*sufR* (Fig. 4). A direct comparison of the expression data confirmed a uniformly reduced expression of the *suf* operon in *Mtb*Δ*sufR* as compared to wt *Mtb* under standard growing conditions and NO stress (Fig. 5A-D). Since SufR is a putative repressor of the *suf* operon [19], the operon’s diminished expression in *Mtb*Δ*sufR* was unexpected. One possibility is that by replacing 345 bp fragment internal to the *sufR* (+308 to +653 bp) with the hygromycin resistance cassette (*lox-hygrgfp-lox*)[23], we might have interrupted the NO599 inducibility of the downstream *suf* genes (*sufBDCSUT*) in *Mtb*Δ*sufR* (Fig. 5A). Consistent with this, the partial transcript of *sufR* originated from the undeleted region (+1 to +307 bp) that is present upstream to the deleted fragment (+308 to +653 bp) retained NO-inducibility in *Mtb*Δ*sufR* (Fig. 5C-D). The basal expression of the *sufBDCSUT* genes in *Mtb*Δ*sufR* is likely due to alternative transcription start site (TSS2) present upstream of *sufB* (Fig. 5A). As expected, *sufR-Comp* expressing a native copy of *sufR* restored the induction of *sufR* in response to NO, whereas rest of the operon remained basally expressed (Fig. 5B-D and Fig. S7). While the *suf* operon was not induced, the expression of pathways requiring Fe-S clusters remained induced in NO-treated *Mtb*Δ*sufR* and *sufR-Comp*. Thus, we anticipate that the heightened demand for Fe-S clusters under NO stress is unlikely to be satisfied in *Mtb*Δ*sufR* or in *sufR-Comp*. The restoration of NO-inducibility of *sufR* but not of the *suf* operon in *sufR-Comp* provides an opportunity to assess the contribution of SufR other than regulating Fe-S cluster homeostasis under NO stress. Altogether, the transcriptomic data indicate that SufR is mainly required to adjust the NO612 responsive expression of the *suf* operon in *Mtb*.

**Figure 5:**
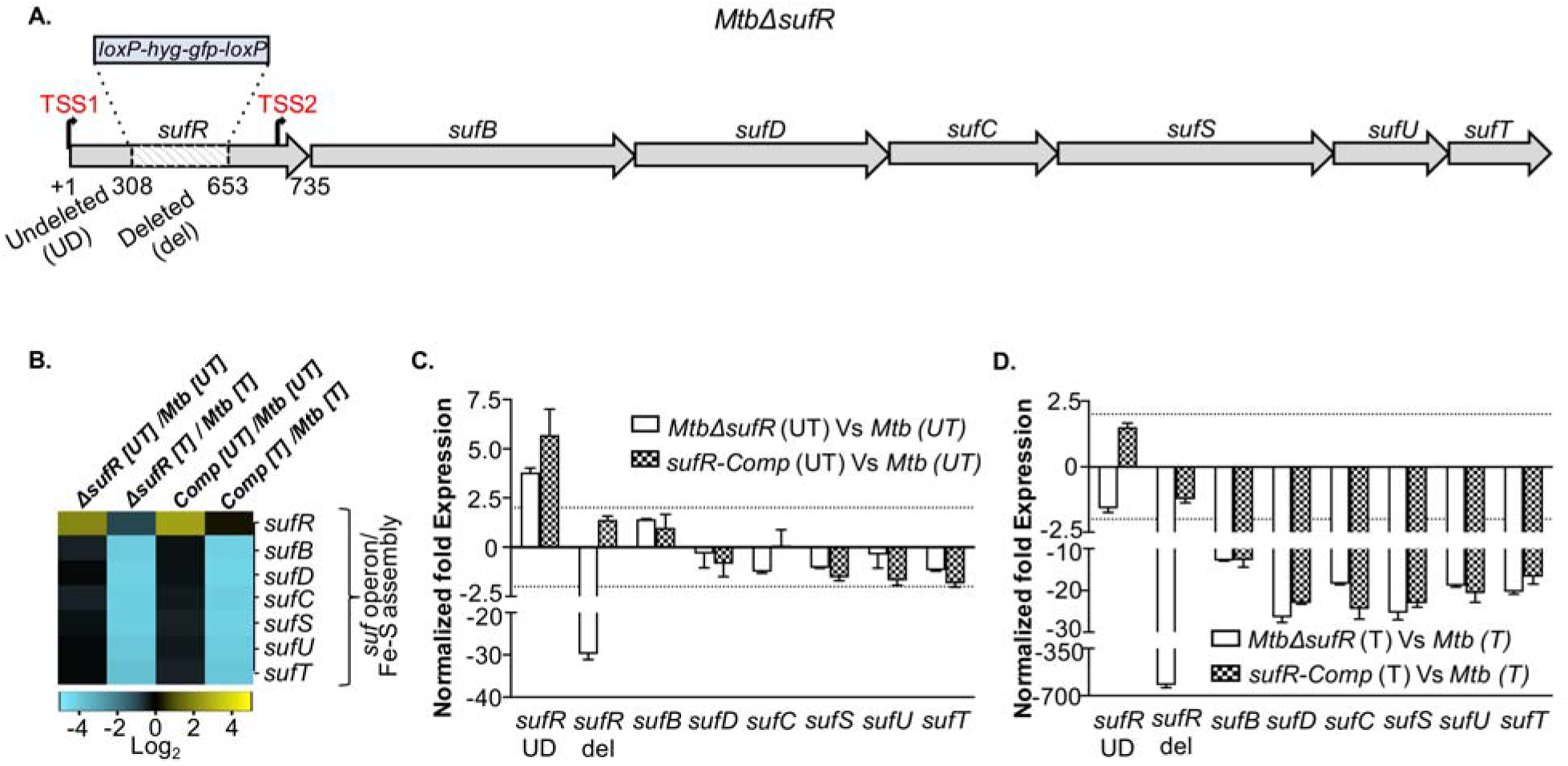
Deregulation of the *suf* operon in *Mtb*Δ *sufR and sufR-comp*. **(A)** Genomic organization of the *suf* operon upon disruption of *sufR*. The internal fragment of *sufR* ORF from +308 bp to +653 bp was deleted (del) and replaced by hygromycin resistance cassette (*loxP-hyg^r-^ gfp-loxP*). The *sufR* sequence from +1 to +307 remained undeleted (UD) and resulted in the expression of an aberrant *sufR* transcript. The two transcription start sites TSS1 (coincides with the start codon ‘*gtg’* of *sufR*) and TSS2 (upstream) of *sufB* were marked as per the published work [19]. **(B)** Exponentially grown cultures of wt *Mtb, Mtb*Δ*sufR, and sufR-comp* were either left untreated (UT) or treated (T) with 500 μM DETA-NO for 4h. Total RNA was isolated and subjected to RNA-seq and qRT-PCR analysis. Heat map depicting expression of the *suf* genes (log_2_fold-change≥1; FDR≤0.05). Note that *sufR* expression detected in *Mtb*Δ*sufR* samples originated from the reads mapping onto the undeleted (UD) region of *sufR*. (**C**, **D**) qRT-PCR data showing the expression of the *suf* genes where *Mtb*Δ*sufR and sufR-comp* were compared with untreated and DETA-NO-treated wt *Mtb*. Results are expressed as mean ± standard deviation (Mean±SD)

### NO irreversibly damages Fe-S clusters of aconitase in *MtbΔsufR*

Altered expression of the Fe-S pathways involved in metabolism and respiration by NO indicates that NO might modulate these processes in *Mtb*. To investigate this idea, we evaluated a 4Fe-4S- containing enzyme aconitase (Acn) activity, which functions as a critical gatekeeper of the TCA cycle, and shows sensitivity to NO due to a solvent-exposed Fe atom [42]. A gradual decrease in Acn activity over time was observed in *Mtb* exposed to 0.5 mM of DETA-NO, indicating Fe-S cluster damage. At 12 h post-exposure, DETA-NO triggered a 40% reduction in Acn activity without decreasing its abundance (Fig. 6A). Importantly, re-culturing DETA-NO-treated *Mtb* in a DETA-NO-free medium significantly restored Acn activity, indicating efficient mobilization of Fe-S cluster regeneration machinery (Fig. 6A).

**Figure 6:**
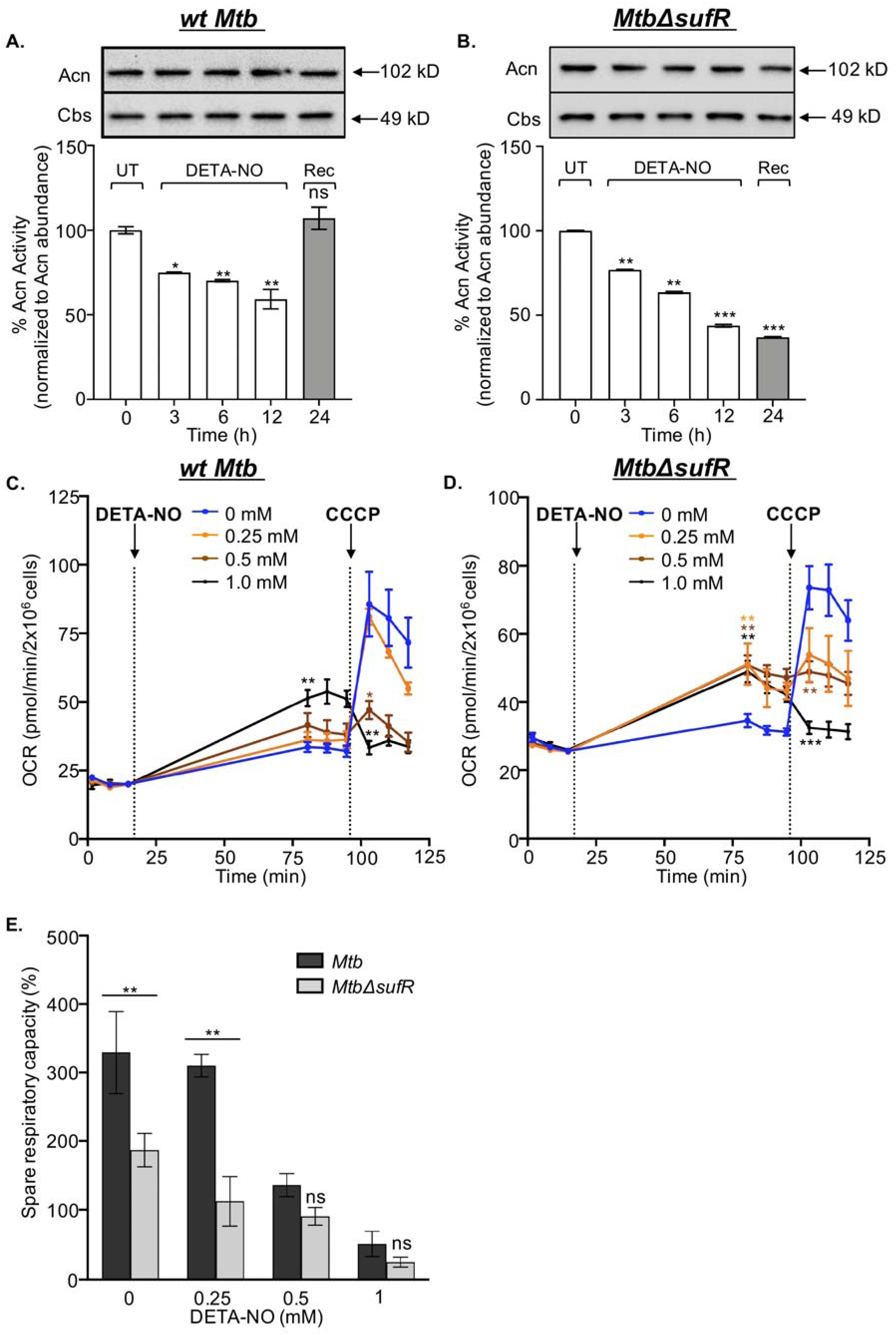
SufR coordinates Fe-S homeostasis, respiratory reserves, and redox balance in response to NO. Aconitase (Acn) activity in response to 0.5 mM DETA-NO. **(A)** *wt Mtb* and **(B)** *Mtb*Δ*sufR.* At 12 h post-treatment, strains were cultured in a DETA-NO free 7H9 broth for 24 h and recovery (Rec) of Acn activity was quantified. Intracellular levels of Acn remained unaltered. A non-Fe-S cluster protein cystathionine-β-synthase (Cbs) of *Mtb* was used as the loading control. A change in oxygen consumption rate (OCR) of (**C**) *wt Mtb* and (**D**) *Mtb*Δ*sufR* was quantified upon addition of indicated concentrations of DETA-NO. The uncoupler CCCP was used to determine the spare respiratory capacity (SRC). The first and second vertical lines indicate point of addition of DETA-NO and CCCP, respectively. **(E)** SRC of *wt Mtb* and *Mtb*Δ*sufR* with increasing concentrations of DETA-NO. Percentage SRC was calculated by subtracting basal OCR (before adding DETA-NO) from CCCP-induced OCR considering basal OCR as 100%. Statistical significance for the OCR was obtained by comparing OCR upon treatment with different doses of DETA-NO with untreated (two-tailed, unpaired Student’s t1131 test.). Comparisons whose P value is <0.05 were indicated with different symbols. Symbols: (*): comparison to 0.25mM; (*): comparison to 0.5mM; and (*): comparison to 1.0 mM */*/* p<0.05; **/**/** p<0.01; ***/***/*** p<0.001. For SRC, one-way analysis of variance (ANOVA) with Bonferroni’s post hoc was used to determine statistical significance. ‘ns’ non1135 significant ‘#’ p<0.05 ‘##’ p<0.01.

We next examined if defective induction of the *suf* operon impairs regeneration of NO-damaged Fe-S clusters in *Mtb*Δ*sufR*. The Acn activity in *Mtb*Δ*sufR* was similar to wt *Mtb* under aerobic growing conditions (Fig. 6B). Moreover, like wt *Mtb*, Acn activity decreased in *Mtb*Δ*sufR* under NO stress over time (Fig. 6B). However, in contrast to wt *Mtb*, reactivation of Acn upon removal of NO stress was absent in *Mtb*Δ*sufR* (Fig. 6B). Importantly, *sufR-Comp* that maintains *sufR* expression but lacks NO-inducibility of the *suf* operon also failed to reinstate Acn activity (Fig. S8A). Data suggest that NO-mediated induction of the *suf* operon rather than SufR alone is critical for the repair of NO-damaged Fe-S clusters in *Mtb*Δ*sufR*. A previous study indicated that Acn activity is dependent upon a stand-alone cysteine desulfurase (IscS) in *Mtb* under standard culture conditions [43]. These findings, along with our data, suggest that *Mtb* prefers IscS under aerobic conditions and Suf system under NO stress for biogenesis of Fe-S clusters. Similar roles were assigned for Isc and Suf systems in *E. coli* [44].

### NO depletes spare respiratory capacity and perturbs redox homeostasis of *MtbΔsufR*

Fe-S cluster-containing enzymes are crucial for maintaining carbon catabolism, oxidative phosphorylation (OXPHOS), and redox balance [36, 45]. Therefore, we exploited Seahorse XF Flux technology to analyze the influence of NO on oxygen consumption rate (OCR) and extracellular acidification rate (ECAR), which are measurable readouts of OXPHOS and glycolysis, respectively [46]. To quantify the basal and maximum rates of OCR and ECAR, we cultured *Mtb* and *Mtb*Δ*sufR* in 7H9-glucose in an XF microchamber, the n exposed it to DETA645 NO, and finally to the uncoupler carbonyl cyanide m-chlorophenyl hydrazine (CCCP). Addition of CCCP stimulates respiration to the maximal capacity manageable by *Mtb*. The difference between basal and CCCP-induced OCR provides an estimate of the spare respiratory capacity (SRC) available for sustaining stress-mediated bioenergetics (*e.g.,* nitrosative and oxidative conditions) [47]. Under normal growing conditions, *Mtb* displayed a basal OCR of 20±0.14 pmoles/min, which increased to 85±11.7 pmoles/min in response to uncoupling stress by CCCP (Fig. 6C). This indicates that *Mtb* normally functions at a submaximal OXPHOS (∼25%) capacity. Under similar conditions, *Mtb*Δ*sufR* utilizes ∼ 35% of its maximal respiratory capacity, which is more than wt *Mtb* (Fig. 6D). Similar to the uncoupler CCCP, NO also depolarizes the cytoplasmic membrane to arrest respiration and growth [48, 49]. Consistent with this, and as seen with CCCP, pretreatment with DETA-NO also increased basal OCR of wt *Mtb* and *Mtb*Δ*sufR* (Fig. 6C, D). However, while 1 mM of DETA-NO was required to increase basal OCR of wt *Mtb* significantly, 0.25 mM was sufficient for *Mtb*Δ*sufR* (Fig. 6C, D). Data suggest that NO stimulated basal OCR, possibly by collapsing proton motive force (PMF), and that *Mtb*Δ*sufR* is more sensitive to membrane depolarization by NO.

We also found that DETA-NO pretreatment progressively reduced the ability of bacteria to increase OCR in response to CCCP (Fig. 6C, D). As a result, DETA-NO significantly decreased SRC in a dose-dependent manner in both *Mtb* and *Mtb*Δ*sufR* (Fig. 6E). However, SRC of *Mtb*Δ*sufR* was significantly lower than wt *Mtb* under normal culture conditions and upon exposure to 0.25 mM DETA-NO (Fig. 6E). These results indicate that *Mtb* mobilizes its reserved respiratory capacity to sustain bioenergetics in response to NO. Data also suggest that the inherently reduced SRC of *Mtb*Δ*sufR* due to diminished Fe-S cluster biogenesis increased its vulnerability towards bioenergetic exhaustion by NO.

Measurement of basal ECAR with and without CCCP treatment indicated that wt *Mtb* and *Mtb*Δ*sufR* operate at a suboptimal glycolytic capacity of 20% and 15%, respectively (Fig. S8B672 C). Similar to OCR, DETA-NO pretreatment progressively reduced the ability of *Mtb* to increase ECAR in response to CCCP (Fig. S8B). However, unlike wt *Mtb*, *Mtb*Δ*sufR* significantly increased basal ECAR and retains CCCP-induced ECAR in response to 0.25 mM and 0.5 mM of DETA-NO (Fig. S8C). This suggests an increased reliance of *Mtb*Δ*sufR* on glycolysis to handle the bioenergetic needs under NO stress. Similar to *Mtb*Δ*sufR*, the NO-induced changes in OCR, SRC, and ECAR were recapitulated in *sufR-Comp* (Fig. S8D-F), indicating that the increased expression of the *suf* operon rather than *sufR* alone is critical for *Mtb’s* response to NO.

Lastly, we asked whether NO perturbed redox homeostasis in *Mtb*. We used a genetic biosensor (Mrx1-roGFP2) to measure the redox potential of a physiologically relevant antioxidant, mycothiol (MSH), as a proxy for the cytoplasmic redox potential (*EMSH*) of *Mtb*[50]. Ratiometric measurements of emission at 510 nm after excitation at 405 and 488 nm can easily quantify any changes in redox physiology [50]. In response to an oxidant or a reductant, the biosensor ratio showed a rapid increase or decrease, respectively [50]. *Mtb* and *Mtb*Δ*sufR* expressing Mrx1- roGFP2 were treated with 0.25 mM, 0.5 mM, 1 mM DETA-NO, and the biosensor ratio was measured. Exposure of *Mtb* to NO did not increase the Mrx1-roGFP2 ratio, indicating that wt *Mtb* robustly maintains cytoplasmic *EMSH* in response to NO (Fig. 7A). In contrast, NO induces a slightly higher oxidative shift in *EMSH* of *Mtb*Δ*sufR* than wt *Mtb* in a dose-dependent manner (Fig. 7A). Using a ROS-sensitive fluorescent dye Cell ROX, we confirmed that NO induces oxidative stress in *Mtb*Δ*sufR* but not in wt *Mtb* (Fig. 7B). Altogether, our data indicate that the *suf* operon’s NO-mediated induction is required to regenerate Fe-S clusters, maintain respiratory reserves, and buffer redox imbalance.

**Figure 7:**
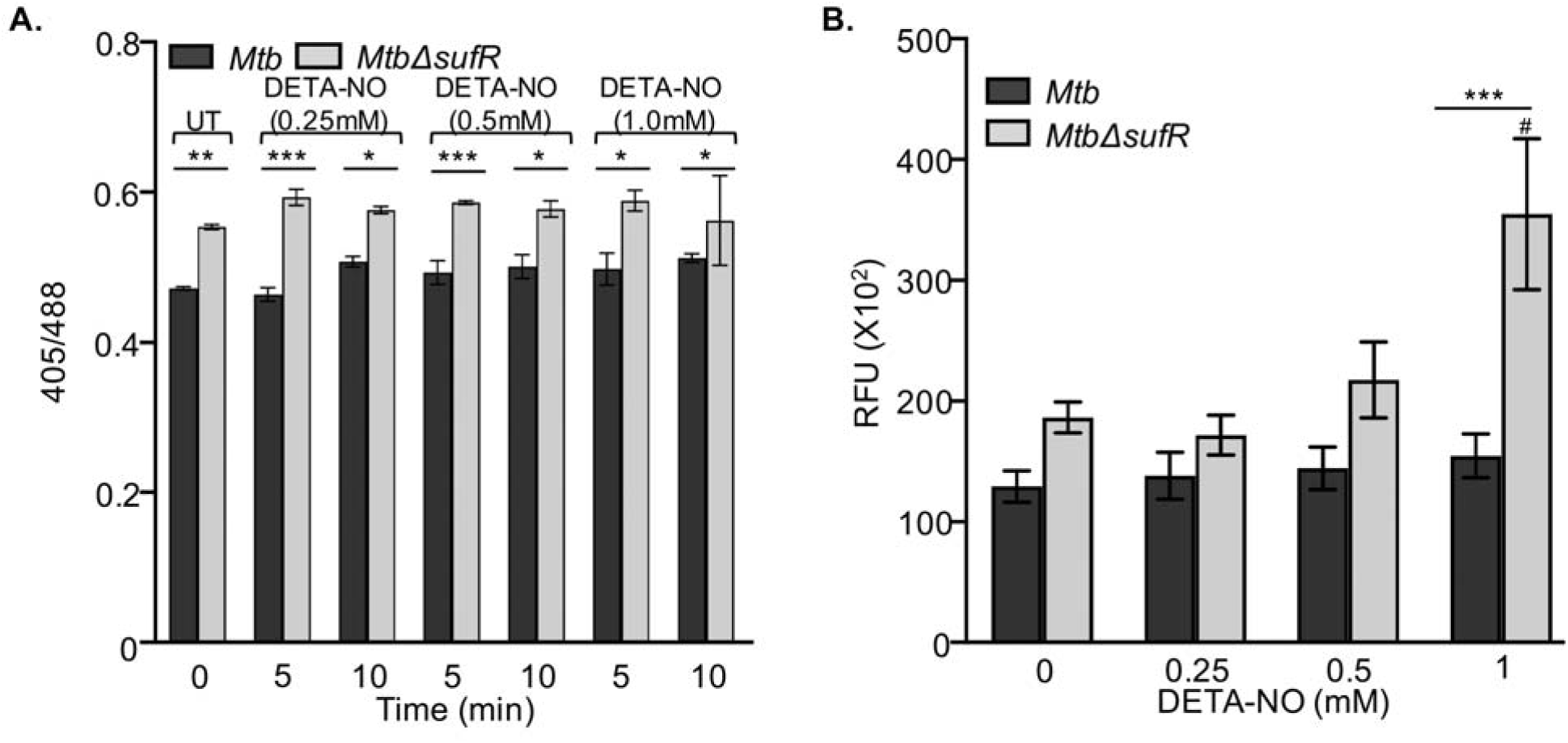
DETA NO induces oxidative stress i n *Mtb*Δ*sufR*. **(A)** wt *Mtb* and *Mtb*Δ*sufR* expressing Mrx1-roGFP2 were exposed to the indicated concentrations of DETA-NO and the biosensor ratio (405/488 nm) was measured using flow cytometry. Data shown are the result of three independent experiments performed in triplicate (mean ± SD). One-way analysis of variance (ANOVA) with Bonferroni’s post hoc test was employed to determine statistical significance. **(B)** wt *Mtb* and *Mtb*Δ*sufR* treated with the indicated concentrations of DETA-NO for 2 h and stained with CellRox Deep Red reagent to measure endogenous ROS. Data shown are the result of three independent experiments performed in triplicate (mean ± SD). Two-way analysis of variance (ANOVA) with Bonferroni’s post hoc test was employed to determine statistical significance between different doses of DETA-NO. Symbols: (*): comparison to *Mtb*, (#): comparison to untreated *Mtb*Δ*sufR*. ‘*’ p<0.01 ‘**’ p<0.01‘***’ p<0.001.

### SufR is required to recover from NO-induced growth arrest and persistence *in vivo*

Next, we investigated the biological consequence of compromised Fe-S homeostasis and bioenergetics by assessing the phenotype of *Mtb*Δ*sufR* under NO stress. First, we investigated the survival phenotype of *Mtb*Δ*sufR* under NO stress *in vitro*. A single dose of various concentrations of DETA-NO (0.5 mM, 1.25 mM, and 2.5 mM) did not influence the survival of *Mtb*Δ*sufR* (**Fig. 8A** and Fig.S9). Repeated exposure to low doses of NO is known to arrest *Mtb*’s growth for an extended duration [51]. Administration of 0.1 mM of DETA-NO every 6 h for 36 h induces an extended period of growth arrest followed by recovery of wt *Mtb* at day 7 post-treatment (**Fig. 8B**). In contrast, *Mtb*Δ*sufR* resumed growth only at day 16 post-treatment with NO (**Fig. 8B**). The *sufR-Comp* strain showed a recovery defect largely similar to *Mtb*Δ*sufR*, reinforcing that the NO inducibility of the entire *suf* operon rather than only *sufR* is necessary for the timely resumption of growth. We confirmed this using CRISPRi-*sufT* strain, which expresses reduced levels of another Fe-S cluster assembly factor-SufT. Similar to *Mtb*Δ*sufR,* diminished levels of *sufT* delayed recovery of *Mtb* from NO-induced growth inhibition (**Fig. 8C**).

**Figure 8:**
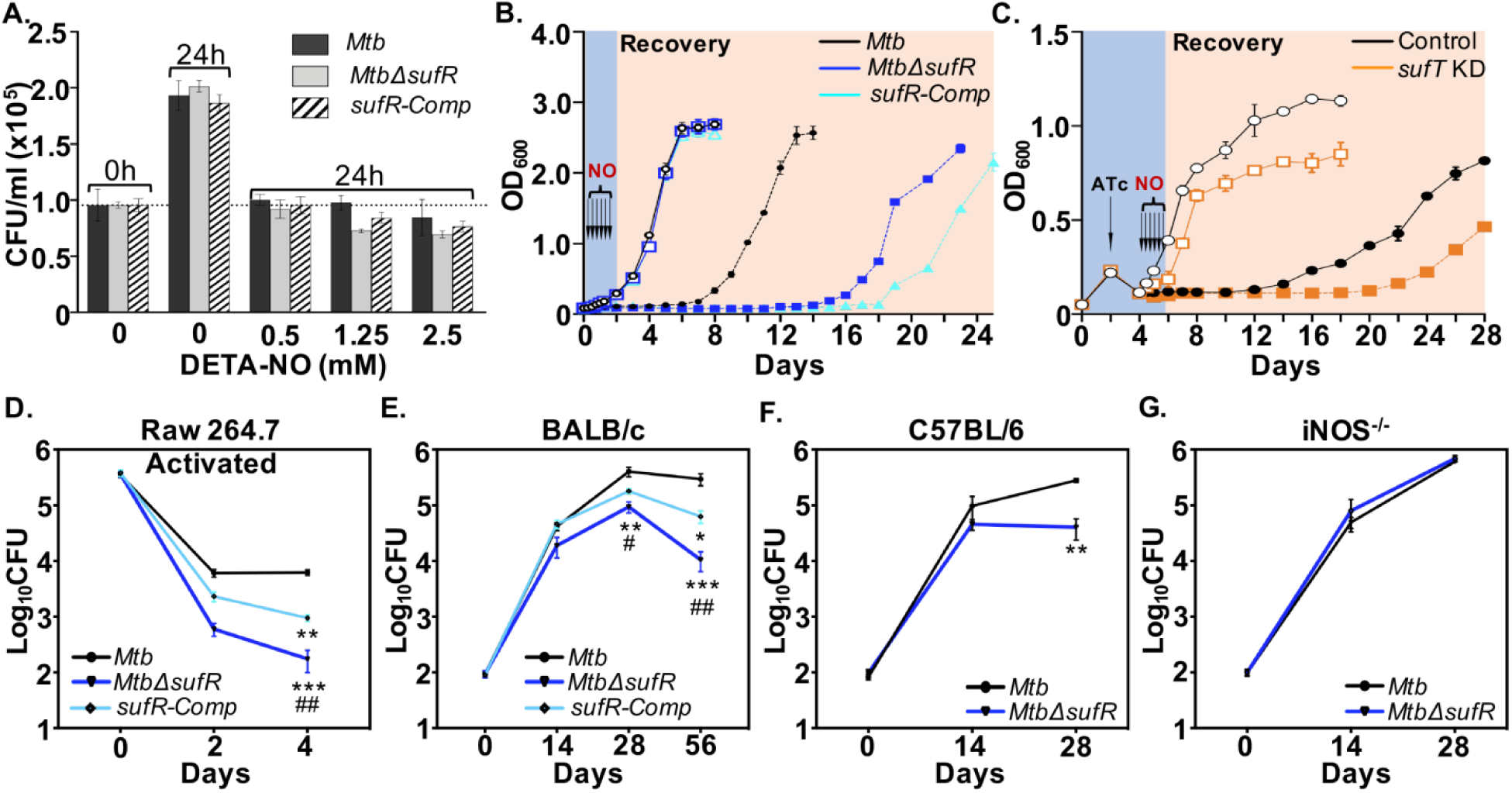
*Mtb*Δ*sufR* displays defect in recovering from NO-induced growth arrest *in vitro* and NO-dependent persistence in mice. **(A)** *Mtb* strains were exposed to DETA-NO for 24 h and survival was monitored by enumerating CFUs. **(B)** *Mtb* strains were either left untreated (UT; empty symbols) or treated (T; filled symbols) with repeated doses of 0.1 mM of DETA-NO for 36 h at an interval of 6 h and growth was monitored over time. **(C)** *Mtb* strains expressing CRISPRi vector without (Control)or with *sufT*-specific guide RNA (sgRNA, *sufT* KD) were left untreated (UT) or treated (T) with 0.1 mM of DETA-NO at an interval of 6 h for 36 h and growth was monitored over time. To deplete SufT, 200 ng/ml of anhydrotetracycline (ATc) was added for the induction of *sufT* specific–guide RNA and dCas9 every 48 h. **(D)** RAW264.7 macrophages activated with IFNγ (100U/mL.) and LPS (100ng/ML.) were infected with *Mtb* strains at a multiplicity of infection (moi) of 10 and survival was monitored by enumerating CFUs. Data are the results of two independent experiments performed in triplicates. Inbred (E) BALB/c mice (n=6), (F) C57BL/6 (n=5), and iNOS knockout (iNOS-/-, n=5) were given an aerosol challenge with *Mtb* strains and assessed for survival in lungs at indicated time points. Results are expressed as Mean±SD. Two-way analysis of variance (ANOVA) with Bonferroni’s post hoc test was employed to determine statistical significance for the intramacrophage and pulmonic load between different strains. p value is <0.05 were indicated with symbols. Symbols: (*): comparison to wt *Mtb*; (#): comparison to *sufR-Comp*. */# p<0.05; **/## p<0.01; *** p<0.001. In fig B and C open and close symbol are untreated and DETA-NO treated respectively.

Immunologically activated macrophages are known to induce nitrosative stress in *Mtb* [52]. In line with this, *Mtb*Δ*sufR* showed 10- and 15-fold reduced survival in immune-activated RAW 264.7 macrophages as compared to wt *Mtb* at day 2 and 4 post-infection, respectively (Fig. 8D). These observations prompted us to investigate the NO-dependent phenotype of *Mtb*Δ*sufR in vivo*. Previous studies have reported the requirement of SufR for persistence of *Mtb* in mice [22, 53]. However, it remains to be addressed if SufR coordinates pathogen’s persistence in response to NO. wt*Mtb* and *Mtb*Δ*sufR* showed comparable growth in the lungs of BALB/c mice during the acute phase of infection (0-2 weeks) (Fig. 8E). However, *Mtb*Δ*sufR* was cleared progressively from the lungs, with a more than 1.5-log decline in bacterial burden by 8 weeks (Fig. 8E). The histopathological changes observed in animals’ lungs at 8 weeks post-infection were proportionate to the bacterial burden (Fig. S10). The magnitude of pulmonary damage was highest in case of wt *Mtb* (2.75±0.5), intermediate in *sufR*-Comp (2.0±0.0), and lowest in *Mtb*Δ*sufR* (0.25±0.50) (Fig. S10B). Lastly, to clarify NO’s role in the persistence defect of *Mtb*Δ*sufR*, we infected a highly susceptible mouse strain lacking inducible nitric oxide (iNOS- /-) [5]. We found that the persistence defect of *Mtb*Δ*sufR* was abolished in iNOS-/- mouse, indicating the requirement of SufR for the persistence of *Mtb* in response to NO (Fig. 8F-G).

Surprisingly, the persistence defect of *Mtb*Δ*sufR* was somewhat rescued in animals and macrophages infected with *sufR-Comp* (Fig. 8D-E). Since *suf* operon’s expression is also responsive to Fe-limitation encountered *in vivo* (41), it is possible that SufR induces the expression of *suf* operon under Fe-limitation from a promoter that is distinct from NO-responsive promoter in *sufR-Comp*. Consistent with this, overexpression of SufR led to its binding inside the ORF of Rv1461 (*sufB*) in *Mtb* [54]. Moreover, several other transcription factors (*e.g., Rv0081*, *Rv0023*, *Rv1189*, *Rv3765c*, *Rv3849*, *Rv0260c*, and *glnR*) bind and alter the expression of *suf* operon [54, 55]. Some of these transcription factors, along with SufR, could regulate *suf* operon’s expression to restore persistence of *sufR-Comp* in animals and macrophages. Alternatively, other enzymes involved in Fe-S cluster coordination in *Mtb* (*e.g.,* IscS) (42) partially counterbalance the repressed Suf system’s effect in *sufR-Comp in vivo*. Consistent with this, our unpublished data suggest that IscS and Suf systems compensate for the loss of each other in mediating *Mtb*’s survival inside macrophages. Altogether, data show that SufR enables NO-dependent persistence of *Mtb* during infection.

## Conclusions

Fe-S cluster production is tightly regulated to promote Fe-S formation when the necessity for the clusters is heightened (*e.g.,* ROI/RNI/iron-limitation) and to limit unnecessary production when the demand is low (e.g., hypoxia) (43). Deregulation of Fe-S cluster biogenesis can lead to toxic accumulation of iron and polysulfides inside cells (43). Therefore, the calibrated expression of Fe-S cluster biogenesis is important. Here we show that *Mtb* SufR is required to regulate Fe-S cluster biogenesis in *Mtb* under NO stress. Our findings provide mechanistic insights into how *Mtb* exploits Fe-S cluster regulation and biogenesis under NO stress to favor the pathogen’s persistence.

We found that NO more severely inhibits spare respiratory capacity of *Mtb*Δ*sufR* as compared to wt *Mtb*. Since spare respiratory capacity depends mainly on the recruitment of previously inactive respiratory complexes (55), the SUF system’s sustained activation can provide a reserve of Fe-S cluster-containing respiratory complexes to maintain electron transfer in response to NO. Induction of the *suf* operon in response to conditions that damage Fe-S clusters (*e.g.,* H2O2, NO, iron-starvation, antibiotics, phagosomal pH, and sputum) [22, 23, 29, 30, 56–59] indicates that *Mtb* relies on Suf-dependent Fe-S cluster coordination to maintain persistence. The capacity of SufR to sense and respond to a range of cues, such as NO, H2O2, and iron limitation, possibly empowers *Mtb* to transduce different redox signals into transcriptional responses crucial for persistence *in vivo*.

One limitation of our study is the lack of complementation in various *in vitro* assays. Since the polar effects of *sufR* disruption interfered with NO-inducibility of the downstream *suf* genes, the phenotypic changes exhibited by *Mtb*Δ*sufR* were mainly due to basal expression of the *suf* operon. Consistent with this, the restoration of expression of *sufR* alone did not rescue the phenotype of *Mtb*Δ*sufR in vitro*. Surprisingly, while both *Mtb*Δ*sufR* and *sufR-Comp* showed defective recovery from NO-mediated growth arrest, the mutant strain recovered earlier than the complemented strain. In this context, we noticed that NO exposure upregulated the DOS dormancy regulon more in *Mtb*Δ*sufR* as compared to wt *Mtb* but below 2-fold (FDR< 0.05) cutoff (Fig. S11). Notably, the expression of the DOS regulon was restored to *wt Mtb* levels in *sufR-Comp* under NO stress (Fig. S11). Therefore, marginally better recovery of *Mtb*Δ*sufR* than *sufR-comp* could be a consequence of elevated DOS regulon in the mutant. Agreeing to this, an *Mtb* strain lacking DOS dormancy regulator (*Mtb*Δ*dosR*) showed defective recovery from NO775 mediated growth cessation [51]. Interestingly, a previous study reported overexpression of the *suf* operon in *Mtb*Δ*dosR* under hypoxia [15], signifying a regulatory loop between SufR and DosR in *Mtb*. In addition to SufR and DosR/S/T system, the Fe-S cluster containing regulators such as WhiB3 and WhiB1 also respond to NO [25, 41, 60]. Further, using bacterial-one-hybrid system, another study reported binding of WhiB3 to the promoter region of *sufR* [55], suggesting that further experiments are needed to fully understand the mechanism underlying the regulation of the *suf* operon in *Mtb*.

Previous studies on the *sufR* mutant did not clarify the polar effects on the downstream *suf* genes. Pandey *et al.*, reported a marginal induction of *sufD*, *sufC*, and *sufT* and a basal expression of *sufB*, *sufS*, and *sufU* in the *sufR*-deleted strain (Δ*SufRTB)* [22]. Δ*SufRTB* grew similar to wt *Mtb* under standard growth conditions but showed survival defect under redox stress and inside macrophages [22]. The mutant also displayed persistence defect in mice [22]. In contrast to our findings, Pandey *et al*., reported a significantly better survival of the *sufR*-complemented strain (Δ*SufRTB::pJEBsufRTB)* than wt *Mtb* under diverse *in vitro* stress conditions and macrophages, and full rescue of the persistence defect in mice [22]. Another study generated three identical truncated mutants of *sufR* (ΔRv1460stop_1.19, ΔRv1460stop_5.19, and ΔRv1460stop_5.20) by introducing a premature stop codon at position 122 [19]. Surprisingly, ΔRv1460stop_1.19 and ΔRv1460stop_5.19 grew slowly than wt *Mtb* under standard growing conditions, whereas growth of ΔRv1460stop_5.20 was comparable to wt *Mtb*. Intriguingly, the activity of Fe-S cluster enzymes was not affected in ΔRv1460stop_5.20 but diminished in the reported *sufR*796 complemented strain [19]. None of these studies examined the expression of full *suf* operon both in the *sufR* mutant and the complemented strain under normal and/or NO stress conditions to rule out polar effects. We believe that the reported discrepancies in the *sufR* complementation could be due to the use of non-native promoters (*e.g.,* mycobacterium optimum promoter [MOP] [22] and *hsp60* [19]) to restore SufR expression in previous studies rather than the NO-responsive native *sufR* promoter used in this study. Altogether, future work is required to explore the breadth of SufR-mediated gene regulation and the role of additional regulators in coordinating the expression of the *suf* operon.

Lastly, the *suf* operon was uniformly induced in clinical isolates of *Mtb* belonging to five globally circulating lineages during survival inside macrophages [61]. These results indicate that regulation of Fe-S cluster biogenesis is a part of core processes that remain conserved in diverse *Mtb* lineages evolved under selection pressure inside the human host. Altogether, we propose a new model of mycobacterial persistence in which SufR senses NO through its Fe-S cluster to coordinate Fe-S cluster biogenesis and regulate metabolism, respiration, and redox balance (Fig. 9).

**Figure 9:**
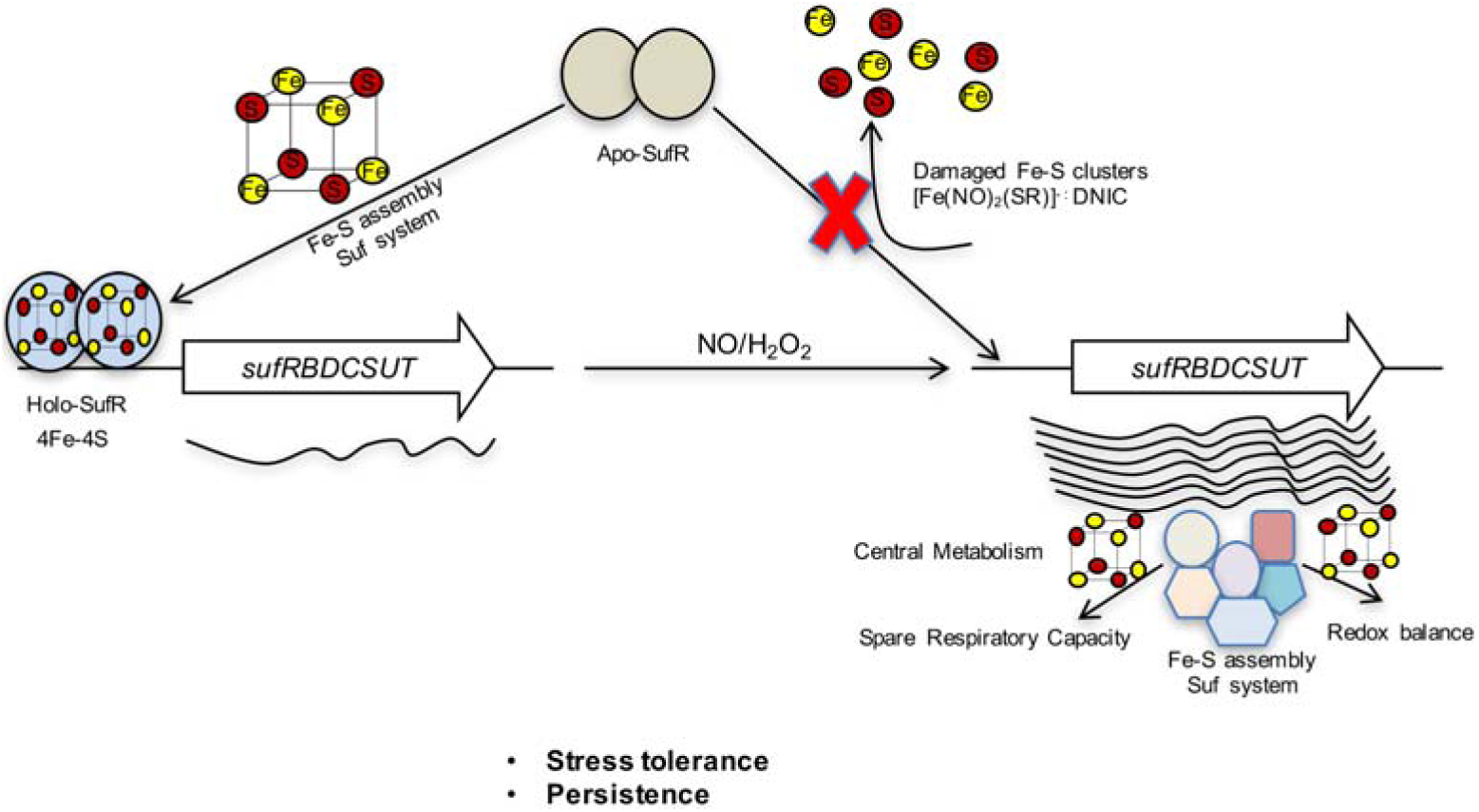
A model for SufR-mediated regulation of Fe-S cluster biogenesis in *Mtb*. SufR appears to coordinate the expression of the *suf* operon in a negative feedback loop. Under Fe-sufficient conditions, the Fe-S cluster bound form of SufR (holo-SufR) binds to the promoter of the *suf* operon and represses Fe-S cluster biogenesis. Exposure to NO or H_2_O_2_ damages Fe-S clusters to generate DNIC or apo-forms of SufR, respectively, which abrogates SufR DNA-binding resulting in de-repression of the *suf* operon. The elevated expression of the SUF Fe-S assembly system regenerates Fe-S clusters of metabolic enzymes, respiratory complexes, and redox sensors to maintain Fe-S cluster homeostasis, spare respiratory capacity, and redox balance. The NO-sensing properties of SufR via its 4Fe-4S cluster and persistence defect exhibited by *Mtb*Δ*sufR* in mice in an iNOS-dependent manner suggest that SufR integrates host environmental signals to Fe-S cluster homeostasis in *Mtb*.

## ASSOCIATED CONTENT

### Supporting Information

Supporting text files containing supplementary figures and supplementary tables information Figure S1-S11

RNA Seq dataset-Table S1A-D (.Xlsx)

List of oligonucleotides used in this study-Table S2 (.Xlsx)

### Author Contributions

KA, AT, and AS participated in the design of the study. KA, AT, KS, NM, AJ, RKJ, RSR, and SNC, carried out the experiments. AR, VN, BG, GN, and ASN contributed to reagents and analyzed the data. KA and AS conceived the study, supervised the project, analyzed the data and drafted the manuscript. All authors read and approved the final manuscript.

### Funding Sources

The *Mtb* work was supported by the following Wellcome Trust/DBT India Alliance Grants, IA/S/16/2/502700 (AS), IA/E/16/1/503017 (KA), and in part by DBT grants BT/PR13522/COE/34/27/2015, BT/PR29098/Med/29/1324/2018, and BT/HRD/NBA/39/07/2018-19 (A.S.), DBT-IISc Partnership Program grant 22-0905-0006-05-987 436, and the Infosys Foundation. AS and KA are senior- and early-career fellows of Wellcome Trust/DBT India Alliance.

### Notes

The funders had no role in study design, data collection and analysis, decision to publish, or preparation of the manuscript. We confirm that no competing financial interests exist.

## ACKNOWLEDGMENT

We are thankful to Awadhesh Pandit and Next Generation Genomics Facility (NGGF) at National Centre for Biological Sciences, Bangalore, for conducting the RNA-sequencing. We thank BSL3 facilities at CIDR, IISc Bangalore. We thank A. Varalakshmi (Sophisticated Analytical Instrument Facility, IIT Chennai, India) for excellent technical help in EPR experiments.

## ABBREVIATIONS

Mtb: Mycobacterium tuberculosis
NO: nitric oxide
iNOS: inducible nitric oxide synthase
DNIC: dinitrosyl-iron dithiol complex
DTH: sodium dithionite
CD: circular dichroism
EMSA: electrophoretic mobility shift assay
DETA-NO: diethylenetriamine-nitric oxide
OCR: Oxygen Consumption Rate
SRC: spare respiratory capacity
CCCP: carbonyl cyanide m-chlorophenyl hydrazine
ECAR: extracellular acidification rate

## REFERENCES

1. Mishra, B.B., et al., Nitric oxide controls the immunopathology of tuberculosis by inhibiting NLRP3 inflammasome-dependent processing of IL-1beta. Nat Immunol, 2013. 14(1): p. 52–60.

2. Voskuil, M.I., et al., Inhibition of respiration by nitric oxide induces a Mycobacterium tuberculosis dormancy program. J Exp Med, 2003. 198(5): p. 705–13.

3. Nathan, C. and M.U. Shiloh, Reactive oxygen and nitrogen intermediates in the relationship between mammalian hosts and microbial pathogens. Proc Natl Acad Sci U S A, 2000. 97(16): p. 8841–8.

4. MacMicking, J., Q.W. Xie, and C. Nathan, Nitric oxide and macrophage function. Annu Rev Immunol, 1997. 15: p. 323–50.

5. MacMicking, J.D., et al., Identification of nitric oxide synthase as a protective locus against tuberculosis. Proc Natl Acad Sci U S A, 1997. 94(10): p. 5243–8.

6. Nicholson, S., et al., Inducible nitric oxide synthase in pulmonary alveolar macrophages from patients with tuberculosis. J Exp Med, 1996. 183(5): p. 2293–302.

7. Mattila, J.T., et al., Microenvironments in tuberculous granulomas are delineated by distinct populations of macrophage subsets and expression of nitric oxide synthase and arginase isoforms. J Immunol, 2013. 191(2): p. 773–84.

8. Wang, C.H., et al., Increased exhaled nitric oxide in active pulmonary tuberculosis due to inducible NO synthase upregulation in alveolar macrophages. Eur Respir J, 1998. 11(4): p. 809–15.

9. Schon, T., et al., Arginine as an adjuvant to chemotherapy improves clinical outcome in active tuberculosis. Eur Respir J, 2003. 21(3): p. 483–8.

10. Schon, T., et al., Effects of a food supplement rich in arginine in patients with smear positive pulmonary tuberculosis--a randomised trial. Tuberculosis (Edinb), 2011. 91(5): p. 370–7.

11. Kumar, A., et al., Mycobacterium tuberculosis DosS is a redox sensor and DosT is a hypoxia sensor. Proc Natl Acad Sci U S A, 2007. 104(28): p. 11568–73.

12. Sardiwal, S., et al., A GAF domain in the hypoxia/NO-inducible Mycobacterium tuberculosis DosS protein binds haem. J Mol Biol, 2005. 353(5): p. 929–36.

13. Cortes, T., et al., Delayed effects of transcriptional responses in Mycobacterium tuberculosis exposed to nitric oxide suggest other mechanisms involved in survival. Sci Rep, 2017. 7(1): p. 8208.

14. Shiloh, M.U., P. Manzanillo, and J.S. Cox, Mycobacterium tuberculosis senses host-derived carbon monoxide during macrophage infection. Cell Host Microbe, 2008. 3(5): p. 323–30.

15. Park, H.D., et al., Rv3133c/dosR is a transcription factor that mediates the hypoxic response of Mycobacterium tuberculosis. Mol Microbiol, 2003. 48(3): p. 833–43.

16. Huet, G., M. Daffe, and I. Saves, Identification of the Mycobacterium tuberculosis SUF machinery as the exclusive mycobacterial system of [Fe-S] cluster assembly: evidence for its implication in the pathogen’s survival. J Bacteriol, 2005. 187(17): p. 6137–46.

17. Sassetti, C.M., D.H. Boyd, and E.J. Rubin, Genes required for mycobacterial growth defined by high density mutagenesis. Mol Microbiol, 2003. 48(1): p. 77–84.

18. Hickok, J.R., et al., Dinitrosyliron complexes are the most abundant nitric oxide-derived cellular adduct: biological parameters of assembly and disappearance. Free Radic Biol Med, 2011. 51(8): p. 1558–66.

19. Willemse, D., et al., Rv1460, a SufR homologue, is a repressor of the suf operon in Mycobacterium tuberculosis. PLoS One, 2018. 13(7): p. e0200145.

20. Coldren, C.D., H.W. Hellinga, and J.P. Caradonna, The rational design and construction of a cuboidal iron-sulfur protein. Proc Natl Acad Sci U S A, 1997. 94(13): p. 6635–40.

21. Nanda, V., et al., Structural principles for computational and de novo design of 4Fe-4S metalloproteins. Biochim Biophys Acta, 2016. 1857(5): p. 531–538.

22. Pandey, M., et al., Iron homeostasis in Mycobacterium tuberculosis is essential for persistence. Sci Rep, 2018. 8(1): p. 17359.

23. Mishra, R., et al., Targeting redox heterogeneity to counteract drug tolerance in replicating Mycobacterium tuberculosis. Sci Transl Med, 2019. 11(518).

24. Crack, J.C., et al., Characterization of [4Fe-4S]-containing and cluster-free forms of Streptomyces WhiD. Biochemistry, 2009. 48(51): p. 12252–64.

25. Singh, A., et al., Mycobacterium tuberculosis WhiB3 responds to O2 and nitric oxide via its [4Fe-4S] cluster and is essential for nutrient starvation survival. Proc Natl Acad Sci U S A, 2007. 104(28): p. 11562–7.

26. Imlay, J.A., Iron-sulphur clusters and the problem with oxygen. Mol Microbiol, 2006. 59(4): p. 1073–82.

27. Freibert, S.A., et al., Biochemical Reconstitution and Spectroscopic Analysis of Iron-Sulfur Proteins. Methods Enzymol, 2018. 599: p. 197–226.

28. Cruz-Ramos, H., et al., NO sensing by FNR: regulation of the Escherichia coli NO- detoxifying flavohaemoglobin, Hmp. EMBO J, 2002. 21(13): p. 3235–44.

29. Voskuil, M.I., et al., The response of mycobacterium tuberculosis to reactive oxygen and nitrogen species. Front Microbiol, 2011. 2: p. 105.

30. Mishra, S., et al., Efficacy of beta-lactam/beta-lactamase inhibitor combination is linked to WhiB4-mediated changes in redox physiology of Mycobacterium tuberculosis. Elife, 2017. 6.

31. Cortes, T., et al., Genome-wide mapping of transcriptional start sites defines an extensive leaderless transcriptome in Mycobacterium tuberculosis. Cell Rep, 2013. 5(4): p. 1121–31.

32. Tamuhla, T., et al., SufT is required for growth of Mycobacterium smegmatis under iron limiting conditions. Microbiology, 2020. 166(3): p. 296–305.

33. Crack, J.C., et al., Iron-sulfur clusters as biological sensors: the chemistry of reactions with molecular oxygen and nitric oxide. Acc Chem Res, 2014. 47(10): p. 3196–205.

34. Beites, T., et al., Plasticity of the Mycobacterium tuberculosis respiratory chain and its impact on tuberculosis drug development. Nat Commun, 2019. 10(1): p. 4970.

35. Baughn, A.D., et al., An anaerobic-type alpha-ketoglutarate ferredoxin oxidoreductase completes the oxidative tricarboxylic acid cycle of Mycobacterium tuberculosis. PLoS Pathog, 2009. 5(11): p. e1000662.

36. Palde, P.B., et al., First-in-Class Inhibitors of Sulfur Metabolism with Bactericidal Activity against Non-Replicating M. tuberculosis. ACS Chem Biol, 2016. 11(1): p. 172–84.

37. Boshoff, H.I., et al., Biosynthesis and recycling of nicotinamide cofactors in mycobacterium tuberculosis. An essential role for NAD in nonreplicating bacilli. J Biol Chem, 2008. 283(28): p. 19329–41.

38. Buchko, G.W., et al., Solution-state NMR structure and biophysical characterization of zinc-substituted rubredoxin B (Rv3250c) from Mycobacterium tuberculosis. Acta Crystallogr Sect F Struct Biol Cryst Commun, 2011. 67(Pt 9): p. 1148–53.

39. Brown, A.C., et al., The nonmevalonate pathway of isoprenoid biosynthesis in Mycobacterium tuberculosis is essential and transcriptionally regulated by Dxs. J Bacteriol, 2010. 192(9): p. 2424–33.

40. Ren, B., et al., Nitric oxide-induced bacteriostasis and modification of iron-sulphur proteins in Escherichia coli. Mol Microbiol, 2008. 70(4): p. 953–64.

41. Kudhair, B.K., et al., Structure of a Wbl protein and implications for NO sensing by M. tuberculosis. Nat Commun, 2017. 8(1): p. 2280.

42. Tortora, V., et al., Mitochondrial aconitase reaction with nitric oxide, S-nitrosoglutathione, and peroxynitrite: mechanisms and relative contributions to aconitase inactivation. Free Radic Biol Med, 2007. 42(7): p. 1075–88.

43. Rybniker, J., et al., The cysteine desulfurase IscS of Mycobacterium tuberculosis is involved in iron-sulfur cluster biogenesis and oxidative stress defence. Biochem J, 2014. 459(3): p. 467–78.

44. Mettert, E.L. and P.J. Kiley, How Is Fe-S Cluster Formation Regulated? Annu Rev Microbiol, 2015. 69: p. 505–26.

45. Lill, R., Function and biogenesis of iron-sulphur proteins. Nature, 2009. 460(7257): p. 831–8.

46. Zhang, J. and Q. Zhang, Using Seahorse Machine to Measure OCR and ECAR in Cancer Cells. Methods Mol Biol, 2019. 1928: p. 353–363.

47. Dranka, B.P., B.G. Hill, and V.M. Darley-Usmar, Mitochondrial reserve capacity in endothelial cells: The impact of nitric oxide and reactive oxygen species. Free Radic Biol Med, 2010. 48(7): p. 905–14.

48. Jones-Carson, J., et al., Nitric oxide disrupts bacterial cytokinesis by poisoning purine metabolism. Sci Adv, 2020. 6(9): p. eaaz0260.

49. Jones-Carson, J., et al., Nitric oxide from IFNgamma-primed macrophages modulates the antimicrobial activity of beta-lactams against the intracellular pathogens Burkholderia pseudomallei and Nontyphoidal Salmonella. PLoS Negl Trop Dis, 2014. 8(8): p. e3079.

50. Bhaskar, A., et al., Reengineering redox sensitive GFP to measure mycothiol redox potential of Mycobacterium tuberculosis during infection. PLoS Pathog, 2014. 10(1): p. e1003902.

51. Leistikow, R.L., et al., The Mycobacterium tuberculosis DosR regulon assists in metabolic homeostasis and enables rapid recovery from nonrespiring dormancy. J Bacteriol, 2010. 192(6): p. 1662–70.

52. MacMicking, J.D., G.A. Taylor, and J.D. McKinney, Immune control of tuberculosis by IFN-gamma-inducible LRG-47. Science, 2003. 302(5645): p. 654–9.

53. Sassetti, C.M. and E.J. Rubin, Genetic requirements for mycobacterial survival during infection. Proc Natl Acad Sci U S A, 2003. 100(22): p. 12989–94.

54. Minch, K.J., et al., The DNA-binding network of Mycobacterium tuberculosis. Nat Commun, 2015. 6: p. 5829.

55. Guo, M., et al., Dissecting transcription regulatory pathways through a new bacterial one-hybrid reporter system. Genome Res, 2009. 19(7): p. 1301–8.

56. Tyagi, P., et al., Mycobacterium tuberculosis has diminished capacity to counteract redox stress induced by elevated levels of endogenous superoxide. Free Radic Biol Med, 2015. 84: p. 344–54.

57. Kurthkoti, K., et al., The Capacity of Mycobacterium tuberculosis To Survive Iron Starvation Might Enable It To Persist in Iron-Deprived Microenvironments of Human Granulomas. mBio, 2017. 8(4).

58. Van den Bossche, A., et al., Transcriptional profiling of a laboratory and clinical Mycobacterium tuberculosis strain suggests respiratory poisoning upon exposure to delamanid. Tuberculosis (Edinb), 2019. 117: p. 18–23.

59. Kumar, M., et al., Identification of Mycobacterium tuberculosis genes preferentially expressed during human infection. Microb Pathog, 2011. 50(1): p. 31–8.

60. Mehta, M. and A. Singh, Mycobacterium tuberculosis WhiB3 maintains redox homeostasis and survival in response to reactive oxygen and nitrogen species. Free Radic Biol Med, 2019. 131: p. 50–58.

61. Homolka, S., et al., Functional genetic diversity among Mycobacterium tuberculosis complex clinical isolates: delineation of conserved core and lineage-specific transcriptomes during intracellular survival. PLoS Pathog, 2010. 6(7): p. e1000988.

62. Gerrick, E.R., et al., Small RNA profiling in Mycobacterium tuberculosis identifies MrsI as necessary for an anticipatory iron sparing response. Proc Natl Acad Sci U S A, 2018. 115(25): p. 6464–6469.

63. Li, H. and R. Durbin, Fast and accurate short read alignment with Burrows-Wheeler transform. Bioinformatics, 2009. 25(14): p. 1754–60.

64. Li, H., et al., The Sequence Alignment/Map format and SAMtools. Bioinformatics, 2009. 25(16): p. 2078–9.

65. Quinlan, A.R. and I.M. Hall, BEDTools: a flexible suite of utilities for comparing genomic features. Bioinformatics, 2010. 26(6): p. 841–2.

66. Robinson, M.D., D.J. McCarthy, and G.K. Smyth, edgeR: a Bioconductor package for differential expression analysis of digital gene expression data. Bioinformatics, 2010. 26(1): p. 139–40.

67. Tian, J., et al., Variant tricarboxylic acid cycle in Mycobacterium tuberculosis: identification of alpha-ketoglutarate decarboxylase. Proc Natl Acad Sci U S A, 2005. 102(30): p. 10670–5.

68. Singh, A.K., et al., Investigating essential gene function in Mycobacterium tuberculosis using an efficient CRISPR interference system. Nucleic Acids Res, 2016. 44(18): p. e143.

69. Crack, J.C., et al., Techniques for the production, isolation, and analysis of iron-sulfur proteins. Methods Mol Biol, 2014. 1122: p. 33–48.

70. Maxam, A.M. and W. Gilbert, Sequencing end-labeled DNA with base-specific chemical cleavages. Methods Enzymol, 1980. 65(1): p. 499–560.

