## Supplemental text for "*Mycobacterium tuberculosis* SufR Responds to Nitric oxide via its 4Fe-4S cluster and Regulates Fe-S cluster Biogenesis for Persistence in Mice"

Correspondence

Amit Singh, Ph.D

Associate Professor

Wellcome Trust-India Alliance Senior Fellow

Department of Microbiology and Cell Biology (MCBL)

Centre for Infectious Disease Research (CIDR)

Indian Institute of Science (IISc)

Bangalore-12

Ph: +91 8022932604

**Supplemental Figures:**

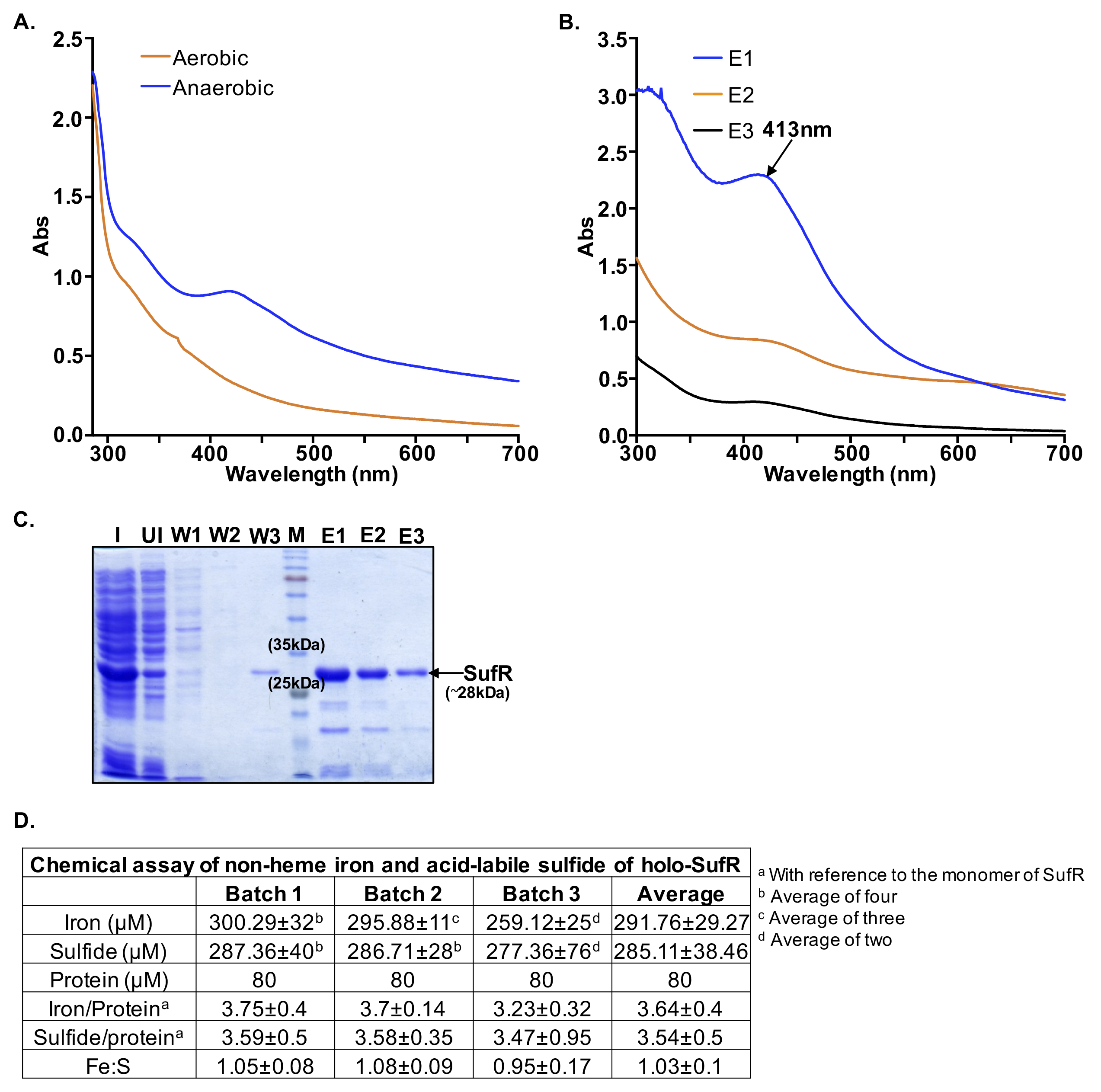

**Figure S1: Purification, spectroscopic characterization and iron-sulfide estimation of *Mtb* SufR (Rv1460):** **(A)** UV-visible spectra of *Mtb* SufR purified under aerobic and anaerobic condition **(B)** UV-visible absorption spectrum of *Mtb* SufR purified in native condition shows the presence of 4Fe-4S cluster indicated by arrow (λmax- 413 nm). Presence of broad peaks at 410-420 nm indicate the presence of a [4Fe-4S] cluster in SufR (Holo-SufR). **(C)** SDS-PAGE confirming the purification of SufR (27.5 kDa). I: Induced, UI: Uninduced, W1, W2, W3: Wash-through fractions, M: Protein marker, E1, E2, and E3: Elution fractions. *Mtb* SufR coordinates 4Fe-4S cluster **(D)** Chemical estimation of Fe and S content in holo-SufR.

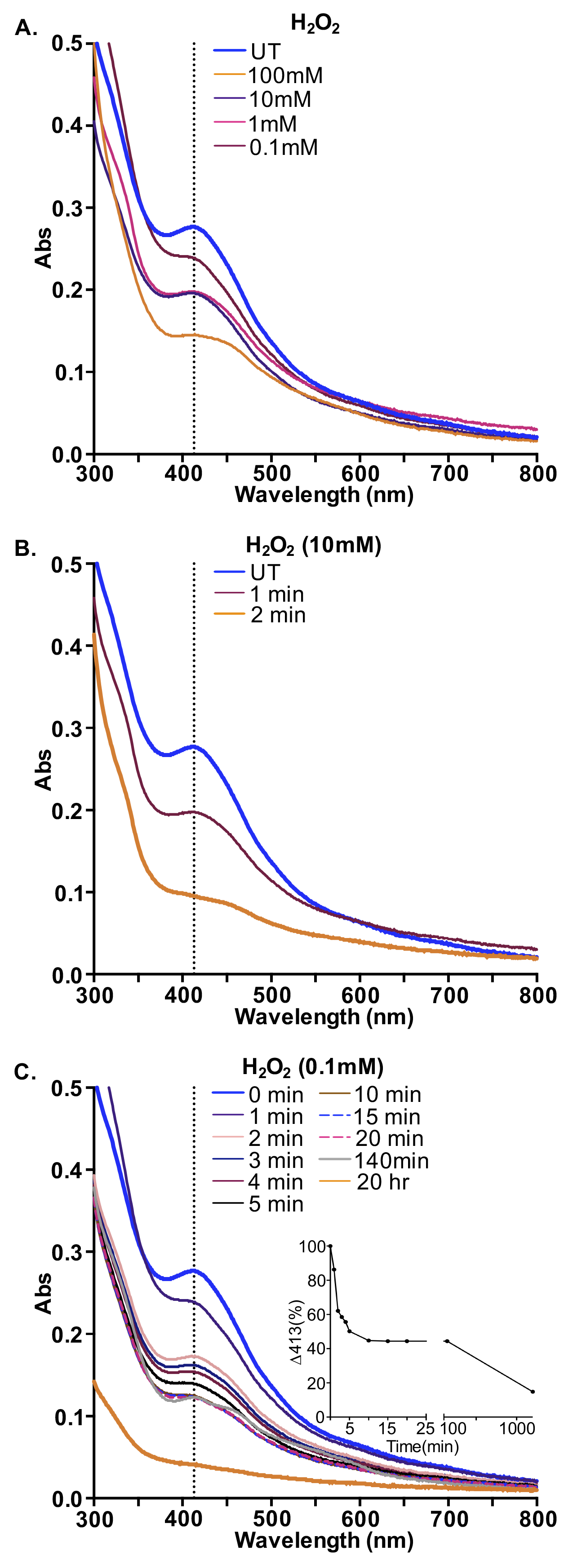

**Figure S2:** ***Mtb* SufR is sensitive to oxidative stress.** Holo SufR was treated either with the **(A)** indicated concentrations of H_2_O_2_ for 1 min or a fixed concentration of H_2_O_2_ **(B)** 10 mM **(C)** 0.1 mM and absorbance was monitored over time. The gradual decrease in absorbance at 413 nm indicates loss of Fe-S cluster. (Inset **C**) Rate of the 4Fe-4S cluster loss was determined by calculating the percent loss of absorbance at 413 nm upon exposure to H_2_O_2_ at various time intervals.

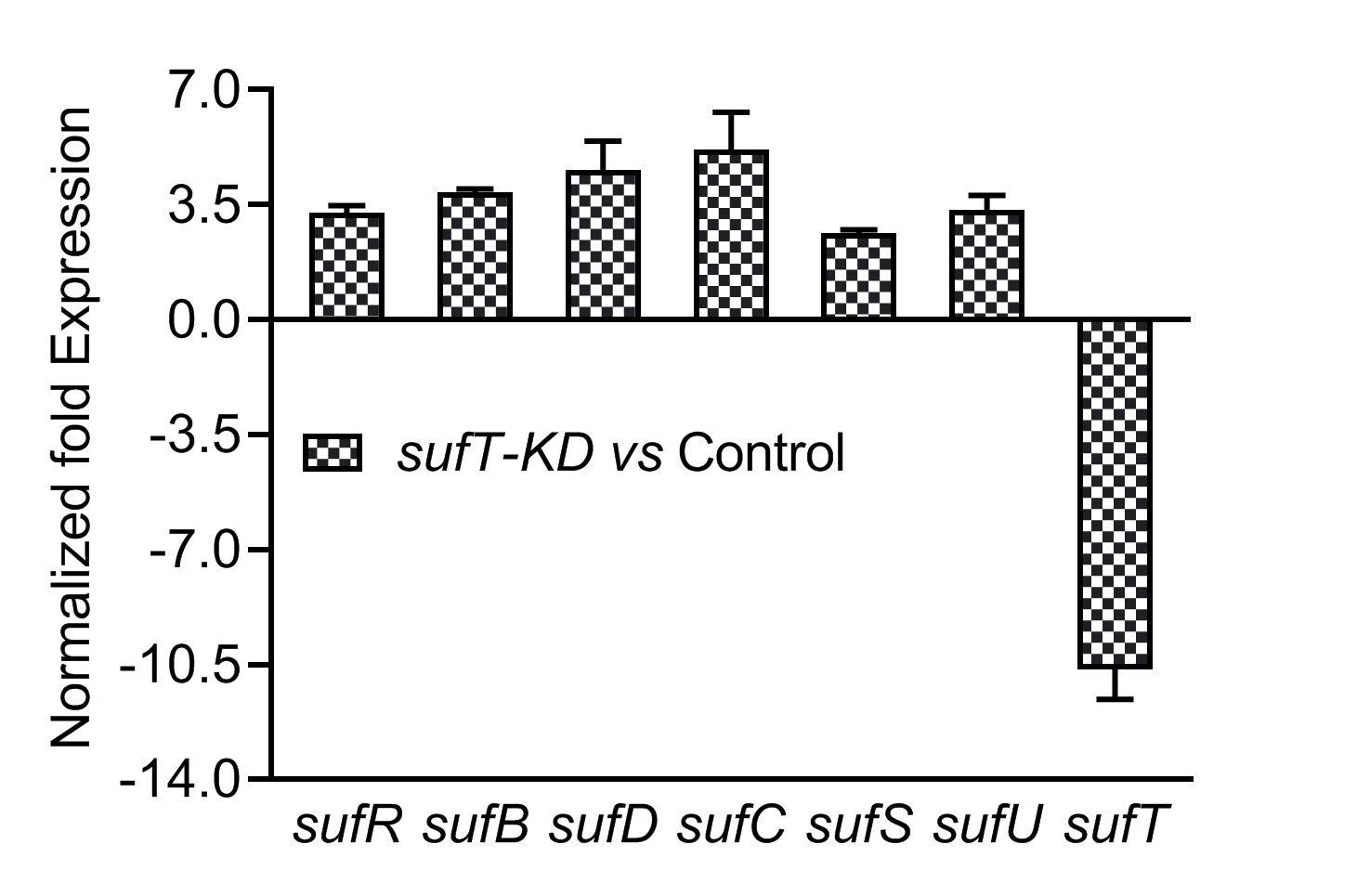

**Figure S3:** **CRISPRi-based knockdown of *sufT* in *Mtb*.** Expression of *sufT* along with *sufR*, *sufB*, *sufD*, *sufC*, *sufS*, and *sufU* in SufT knockdown (KD) strain added with ATc (induced) for 24 h, with respect to control strain not added with ATc (without induction). Data shown are the result of three independent experiments performed in duplicate. Results are expressed as mean±SD.

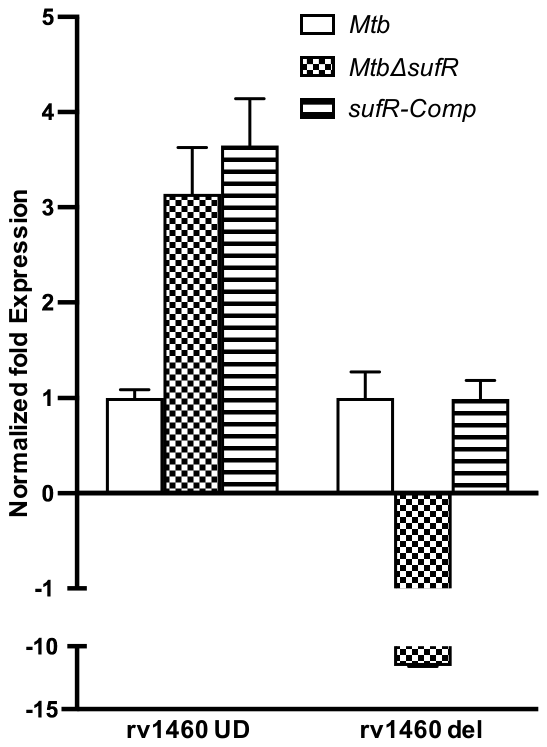

**Figure S4:** **Expression of *sufR*.** *wt Mtb*, *MtbΔsufR* and *sufR*-*Comp* strains were grown and exposed to 500 μM DETA-NO for 4h. Total RNA was isolated and expression of *sufR* was checked using qRT-PCR in the deleted (Rv1460 del) as well as undeleted/intact region (Rv1460 UD) of *sufR* (for deleted and undeleted region please refer **Fig. 5D**). Data shown are the result of three independent experiments performed in triplicate. Results are expressed as mean±SD.

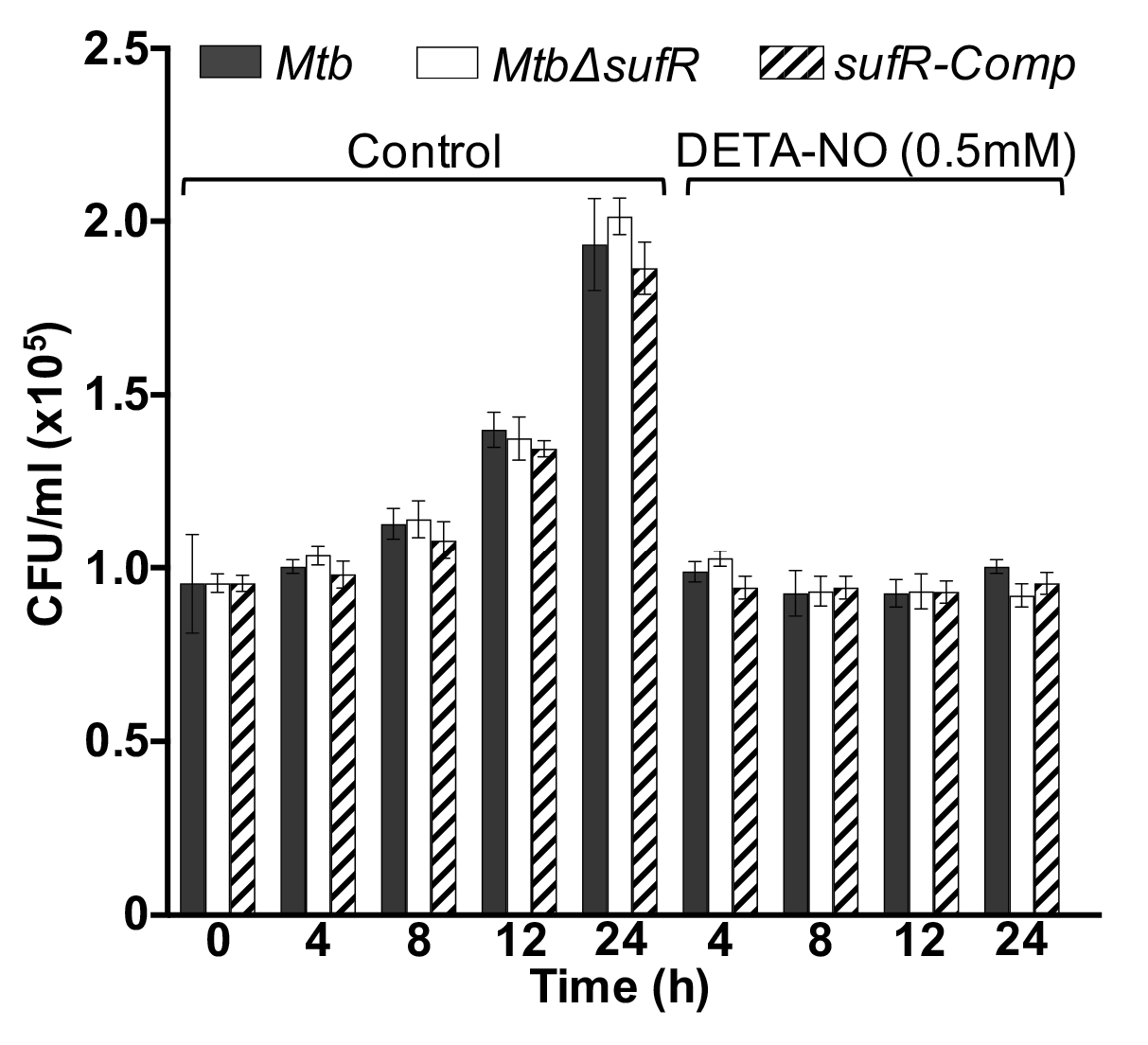

**Figure S5:** **NO arrests growth of *wt Mtb*, *MtbΔsufR* and *sufR–Comp*.** *Mtb* strains were treated with DETA-NO (0.5 mM) and survival was monitored by enumerating CFUs at the indicated time points. Data shown are the result of three independent experiments performed in duplicate. Results are expressed as mean±SD.

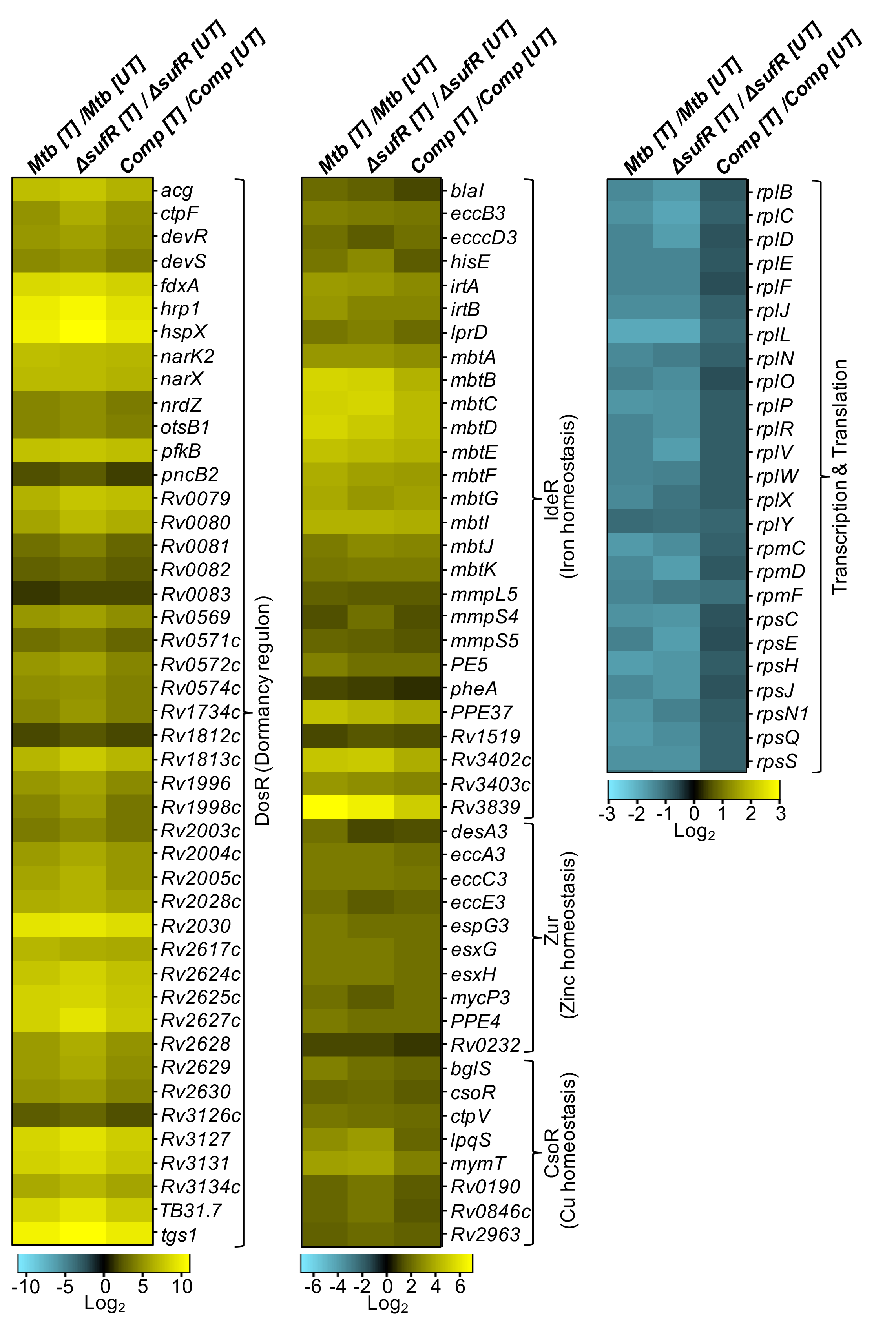

**Figure S6: Regulation of multiple regulons and impact of SufR on the transcriptome of *Mtb* in response to NO stress in *Mtb*.** *wt Mtb*, *MtbΔsufR* and *sufR*-*Comp* strains were grown and exposed to 500 μM DETA-NO for 4h. Total RNA was isolated and subjected to RNA-seq analysis*.* Heat maps depicting expression of genes (log_2_fold-change≥1; FDR≤0.05) coordinating dormancy regulon, Fe-homeostasis, Zn-homeostasis, Cu-homeostasis, transcription and translation for untreated (UT) and 4h of DETA-NO-treated (T) strains from three biological samples.

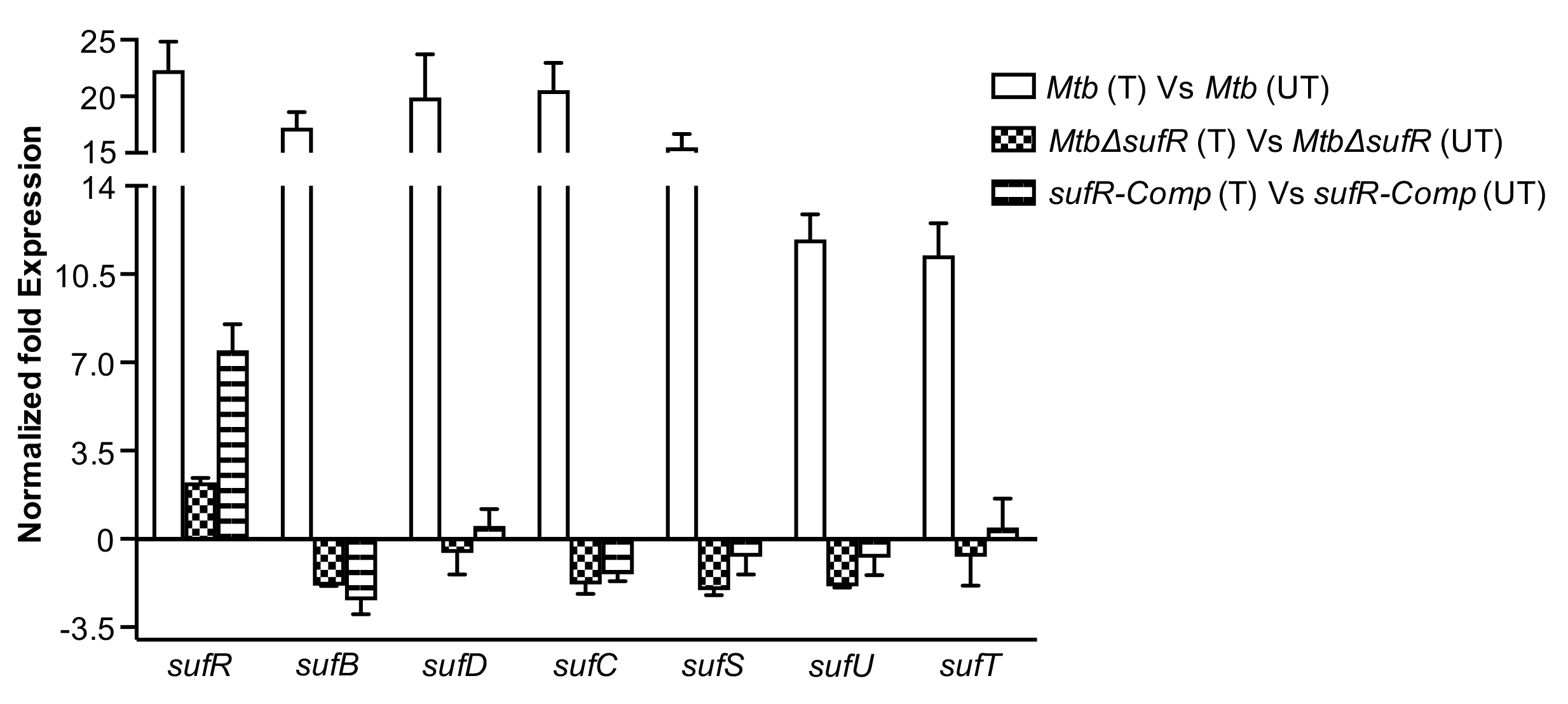
**Figure S7. Expression of *suf* operon upon DETA-NO treatment.** Total RNA from three biological replicates of untreated (UT) and DETA-NO treated (T) *Mtb*, *MtbΔsufR,* and *sufR-Comp* subjected to qRT-PCR. Expression of *suf* operon genes (*sufR*, *sufB*, *sufD*, *sufC*, *sufS*, *sufU* and *sufT*) were compared with untreated counterpart. Results are expressed as mean±SD.

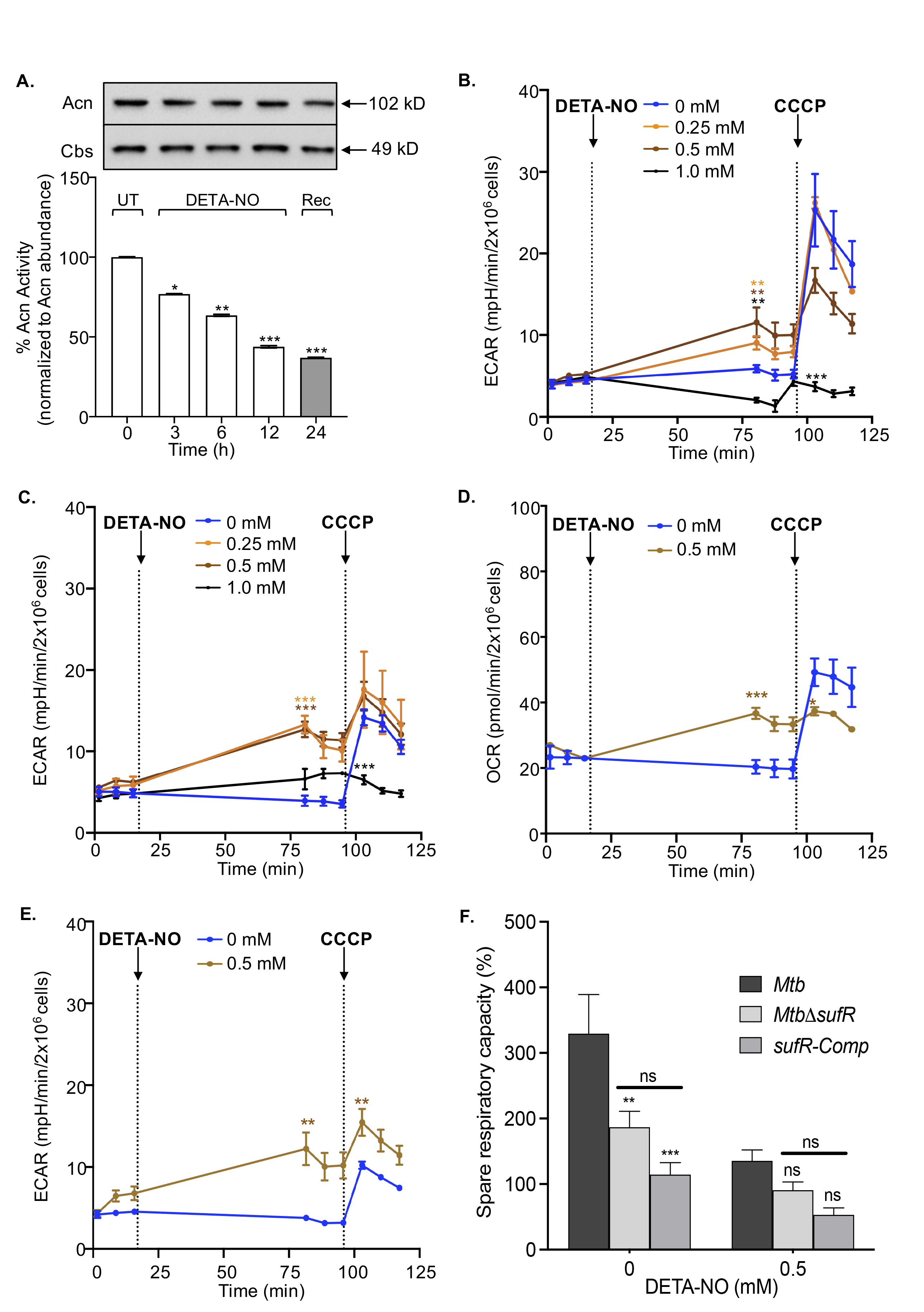

**Figure S8. (A) Aconitase activity of *sufR-Comp* strain upon NO stress.** Aconitase (Acn) activity in *sufR-Comp* upon treatment with 0.5 mM DETA-NO over time. After 12 h, cells were freshly cultured in NO-free 7H9 broth for 24 h and subjected to Acn activity. No recovery in Acn activity was observed after 24 h. The change in enzymatic activity is independent of Acn abundance. Level of Cbs was measured as housekeeping control. Data shown are the result of three independent experiments. Results are expressed as mean±SD. One-way analysis of variance (ANOVA) with Bonferroni's post hoc test was employed to determine statistical significance. Statistical significance was obtained by comparing DETA-NO treated samples at different time points with untreated control (0 h). ‘*’ p<0.05. ‘***’ p<0.001. Extracellular acidification rate (ECAR) of DETA-NO treated (**B**) *wt Mtb* **(C)** *MtbΔsufR* and (**E**)*sufR-Comp*. **(F)** SRC of wt *Mtb* and *MtbΔsufR* and *sufR-Comp* were compared upon 0.5mM concentrations of DETA-NO treatment. Percentage SRC was calculated by subtracting basal OCR (before adding DETA-NO) from CCCP-induced OCR considering basal OCR as 100%. One-way analysis of variance (ANOVA) with Bonferroni's post hoc determined statistical significance. ‘**’ p<0.01 ‘***’ p<0.001. Statistical significance for the ECAR and OCR was obtained by comparing different doses of DETA-NO with untreated (Two-tailed, unpaired Student’s t- test.), Comparisons whose P value is <0.05 were indicated with different symbols. Symbols: (*): comparison to 0.25mM; (*): comparison to 0.5mM; and (*): comparison to 1.0 mM. ‘ns’ non-significant; */*/* p<0.05; **/**/** p<0.01; ***/***/*** p<0.001.

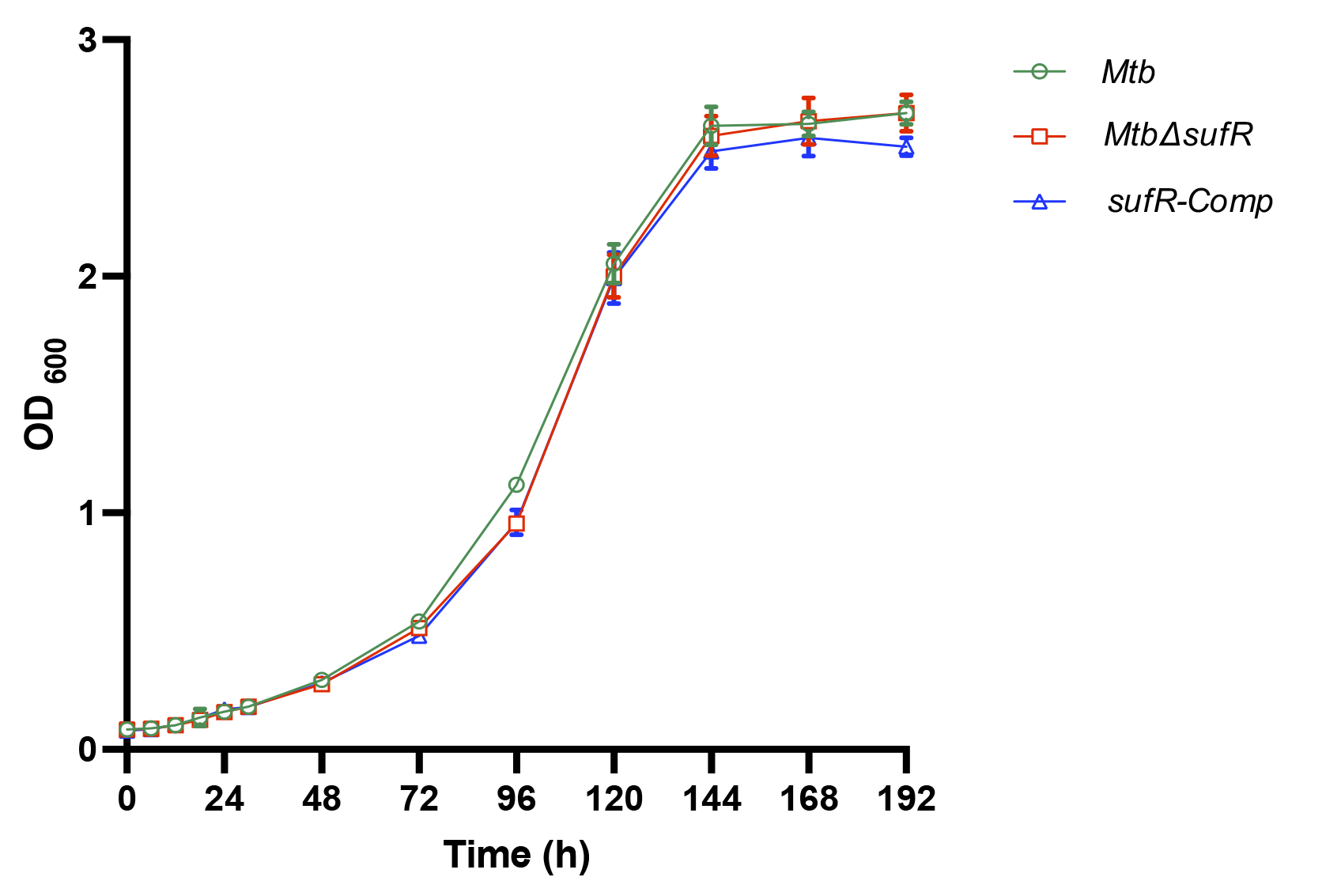

**Figure S9: *MtbΔsufR* does not show growth defect under standard growth condition:** wt *Mtb*, *MtbΔsufR* and *ΔsufR-Comp* strains were cultured under standard growth conditions in 7H9. OD_600_ was monitored at indicated time points. The results shown are the mean and standard deviation of three experiments.

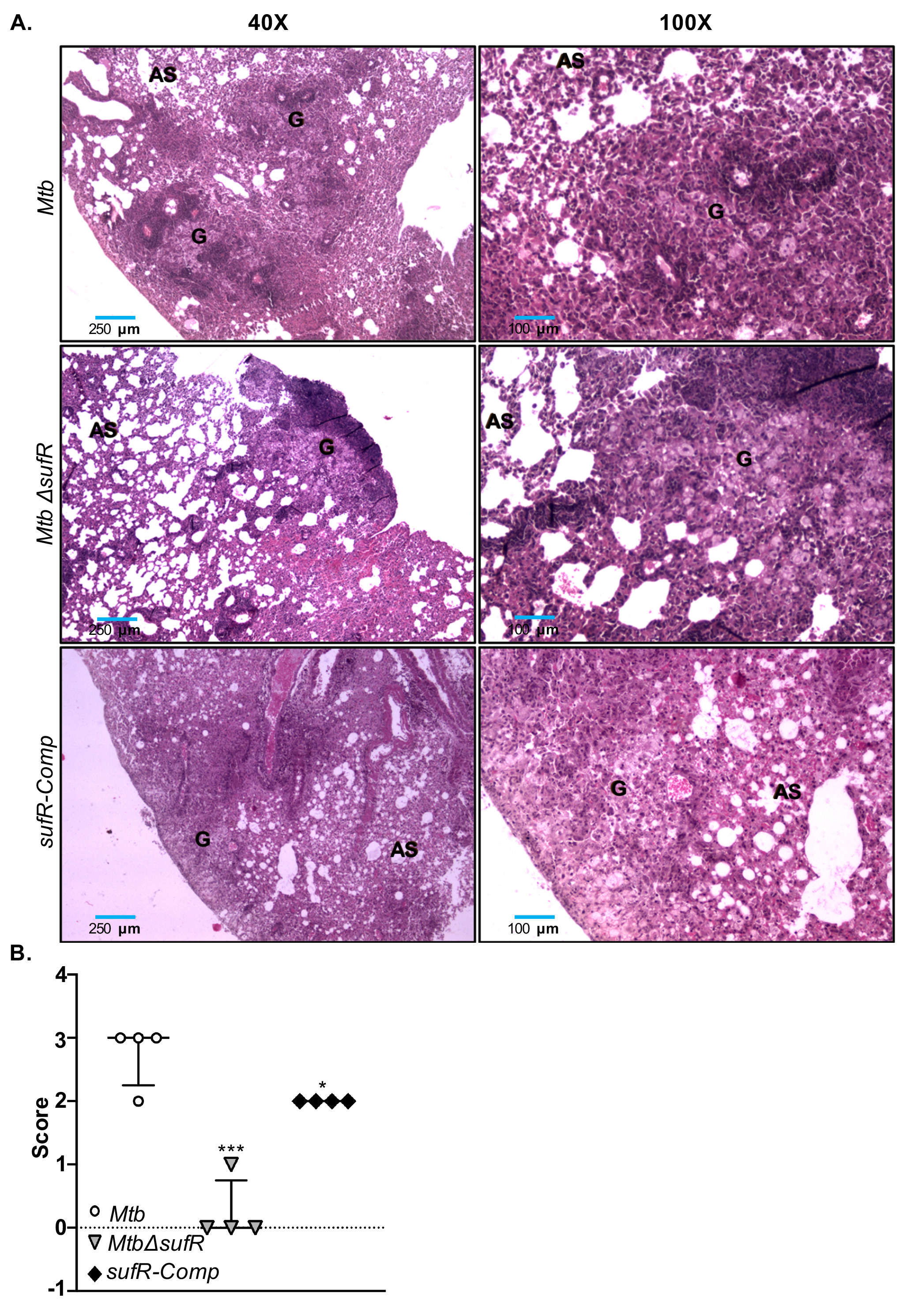

**Figure S10: Histopathology of BALB/c lungs infected with *Mtb*, *MtbΔsufR* and *ΔsufR-Comp* strains.** (**A)** Representative H&E staining of the lung tissue sections of BALB/c mice 8-weeks’ post infection. (**B)** Scatter plots indicate modified granuloma score for lung section histopathology of chronically-infected BALB/c mice across experimental groups at 8-weeks post-infection. Results are expressed as median±inter-quartile range. ‘*’ p<0.05. ‘***’ p<0.001. One-way analysis of variance (ANOVA) with Bonferroni's post hoc test was employed to determine statistical significance. Statistical significance was obtained by comparing with wt *Mtb* (n=4).

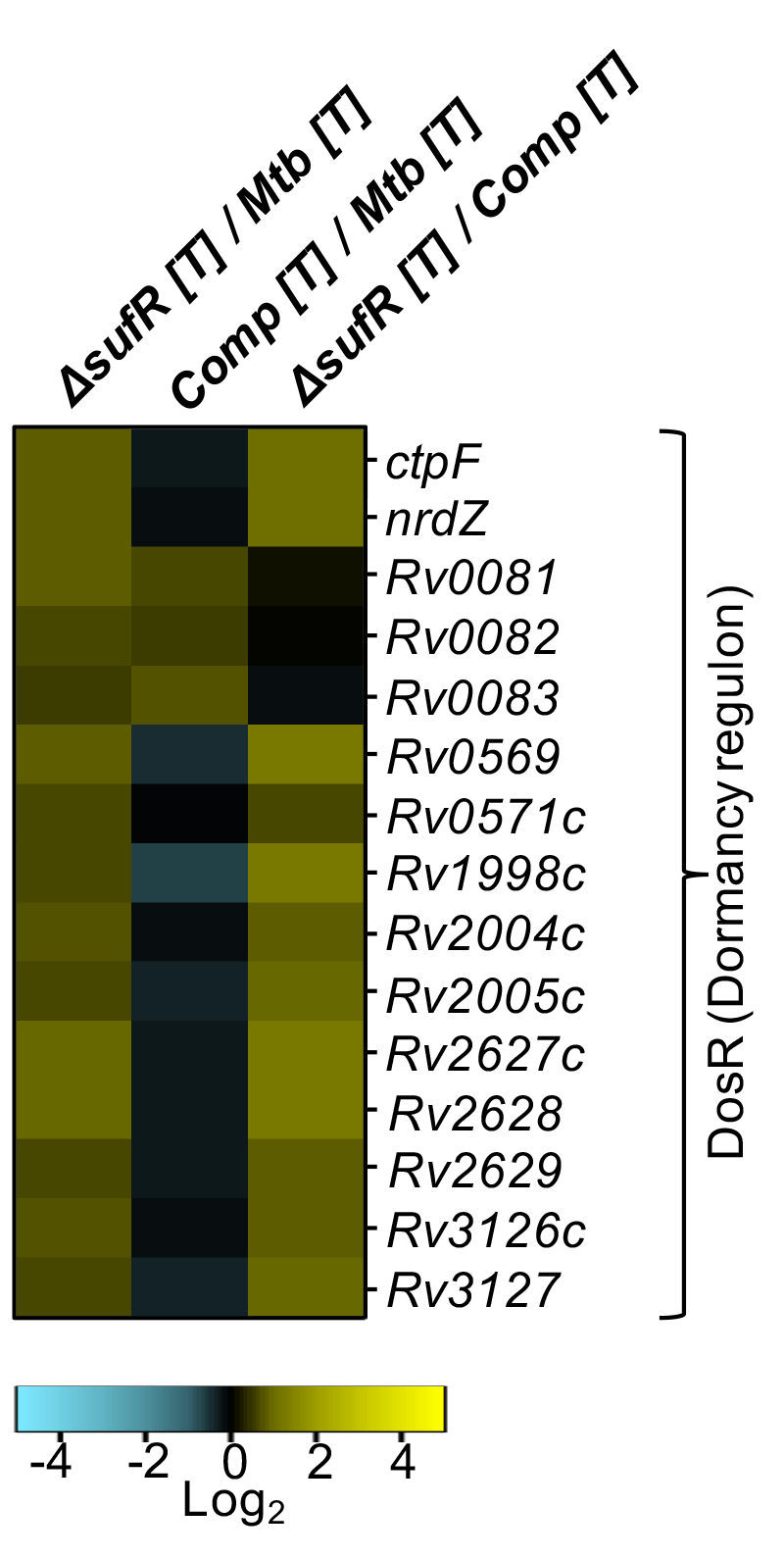

**Figure S11:** **SufR regulates DOS Dormancy Regulon in response to NO.** Total RNA was isolated from three biological replicates of untreated (UT) and DETA-NO treated (T) *MtbΔsufR* and *sufR-Comp* and wt *Mtb* subjected to RNA-seq analysis. Heat maps indicate log_2_ fold changes of differentially expressed genes (DEGs) belonging to various functional categories (obtained from Mycobrowser, EPFL, Lausanne). Genes were considered differentially expressed on the basis of the false discovery rate (FDR) of ≤0.05 and absolute fold change of ≥1.5. “T” and “UT” indicate DETA-NO treated and untreated conditions, respectively.

**Supplemental table**

**List of Supplementary Table:**

**Supplementary Table S1A:** Differentially regulated genes in *wt* *Mtb* upon exposure to 0.5mM DETA-NO as compare to untreated *wt* *Mtb* [log_2_fold-change≥1; FDR≤0.05]. (Enclosed as a separate excel spread sheet).

**Supplementary Table S1B:** Genes differentially regulated in DETA-NO treated *MtbΔsufR* as compared to untreated *MtbΔsufR* [log_2_ fold-change≥1; FDR≤0.05]. (Enclosed as a separate excel spread sheet).

**Supplementary Table S1C:** Genes differentially regulated in DETA-NO treated *Mtb sufR-Comp* as compared to untreated *Mtb sufR-Comp* [log_2_ fold-change≥1; FDR≤0.05]. (Enclosed as a separate excel spread sheet).

**Supplementary Table S1D:** List of Fe-S cluster protein differentially regulated in DETA-NO treated *wt Mtb, MtbΔsufR and sufR-Comp* as compared to untreated wt *Mtb* counterpart. [fold change ≥ 1.5; FDR ≤ 0.05] (Enclosed as a separate excel spread sheet).

**Supplementary Table S2:** Sequences of primers used in this study.
